# Engineering modular cargo loading strategies for carboxysome-derived protein particles

**DOI:** 10.64898/2026.04.24.720684

**Authors:** Claudia A. Mak, Rachelle M. Baumann, Anthony G. Vecchiarelli

**Affiliations:** Department of Biological Chemistry, University of Michigan Medicine, Ann Arbor, Michigan 48109, United States; Department of Molecular, Cellular, and Developmental Biology, University of Michigan, Ann Arbor, MI, 48109, USA

## Abstract

<u>B</u>acterial <u>m</u>icro<u>c</u>ompartments (BMCs) are a diverse and widespread class of protein-based organelle consisting of a semi-permeable protein shell encapsulating an enzymatic core. Along with their native assembly pathway, isolated BMC shell proteins have been shown to assemble into alternative superstructures such as flat sheets and nanotubes. The self-assembly and modularity of BMC shell proteins make them of great interest as modular platforms for applications involving scaffolding, immobilization and compartmentalization. While the assembly of BMC shell proteins into higher-order structures has been well-studied, the design of controllable and modular cargo loading is underdeveloped in comparison. Recently, we reported the pH-controlled assembly of CcmK2 - the major hexameric shell protein of the β-carboxysome BMC - into monodisperse mesh-like microscale particles. Here, we develop a suite of encapsulation strategies for stochastic or targeted loading of various cargos, as well as the direct conjugation of cargo to CcmK2 particles. Our systematic analysis demonstrates that cargo loading and particle assembly can be modulated by the choice of recruitment strategy and the order of cargo introduction. Our findings also reveal a cooperative cargo loading mechanism during assembly that influences particle sizing and apparent morphology. Our study serves as a blueprint for the rational design of tunable cargo loading into engineered BMC-derived microcompartment systems for diverse biotechnological applications.

## INTRODUCTION

Subcellular compartmentalization is a fundamental strategy for spatially organizing metabolic processes and enhancing reaction efficiency. Bacteria have evolved proteinaceous organelles, termed <u>b</u>acterial <u>m</u>icro<u>c</u>ompartments (BMCs), to confine and increase the efficiency of key enzymes within a semi-permeable protein shell.^1^ Hundreds of copies of protein oligomers - hexamers, trimers, and pentamers - tile together to form the complex polyhedral BMC shell.^2,3^ Because BMCs are entirely protein-based and self-assemble, there is great interest in bioengineering BMCs as tunable, modular platforms for programmable cargo encapsulation and the controlled organization of biochemical processes.^4^ Although a major focus of BMC-engineering involves repurposing whole BMCs as nanofactories^5–7^, the complexity of BMC shells and incomplete knowledge of their native assembly - especially how shell proteins encapsulate the enzymatic core - hinder these current efforts. Alternatively, several studies have demonstrated that when isolated, individual BMC shell proteins can assemble into alternative superstructures, such as flat sheets and nanotubes.^8–13^ These superstructures have successfully been engineered into enzymatic scaffolds.^14–16^ The greater utility of BMC shell proteins as modular platforms for bioengineering hinges upon the expansion of available BMC-based architectures and their development into novel functional compartments and scaffolds.

Recently, we demonstrated the assembly of monodisperse spheroidal microscale particles using only CcmK2, the major hexameric shell protein of the β-carboxysome BMC from *Synechococcus elongatus* PCC 7942.^17^ We found that CcmK2 hexamers tile together to form nanotubes, which subsequently cluster into higher order mesh-like particles. The monodispersity, pH-regulated sizing and reversible assembly, high stability, and hierarchical structure of CcmK2 particles situate them as a unique platform for a variety of biotechnological applications, such as enzyme scaffolding for biocatalysis, drug delivery, and bioremediation.^14,18–20^ Future applications of CcmK2 particles, however, necessitate the development of efficient and tunable loading of diverse cargos, such as small molecules and proteins.

Here, we developed multiple approaches for the encapsulation of various cargos by CcmK2 particles: (1) direct conjugation to the CcmK2 hexamer; (2) non-specific, stochastic loading; (3) targeted loading via the fusion of the native carboxysome encapsulation peptide (EP) to a cargo of interest^21^; and (4) incorporation of the SpyCatcher/SpyTag system for covalent anchoring^22,23^. We used these strategies to demonstrate the ability of CcmK2 particles to encapsulate small molecules (fluorescent dyes) and small proteins (mCherry) for proof-of-concept. We identified the length of the linker between the EP and protein cargo to be a modulator of loading capacity and particle assembly. Lastly, we found that the order of cargo introduction dictates the loading mechanism. When cargo is present during particle assembly, loading is governed by a cooperative mechanism in which the cargo increases particle size, thereby enhancing loading capacity. When cargo is introduced to preassembled particles, loading proceeds stochastically, likely via diffusion-limited permeation into particles. Altogether, our findings provide groundwork for the rational design of efficient and tunable cargo loading for diverse biotechnological applications.

## RESULTS

### Small dye molecules can be directly conjugated to CcmK2 for particle loading

Many protein-based nano- and micro-particles encapsulate small molecules and dyes for applications such as drug delivery, therapeutics, and bioimaging.^24–26^ In our previous work, we labeled CcmK2 with Alexa Fluor 488 (MW= 721 Da; CcmK2^488^) and Alexa Fluor 647 (CcmK2^647^) via maleimide conjugation to CcmK2[A68C], which has a strategic alanine to cysteine substitution in a flexible loop region (Figure 1A).^17^ We decorated CcmK2 particles with 0.1% CcmK2^488^ or CcmK2^647^ to ensure sufficient labeling density for imaging without perturbing particle assembly.

**Figure 1.**
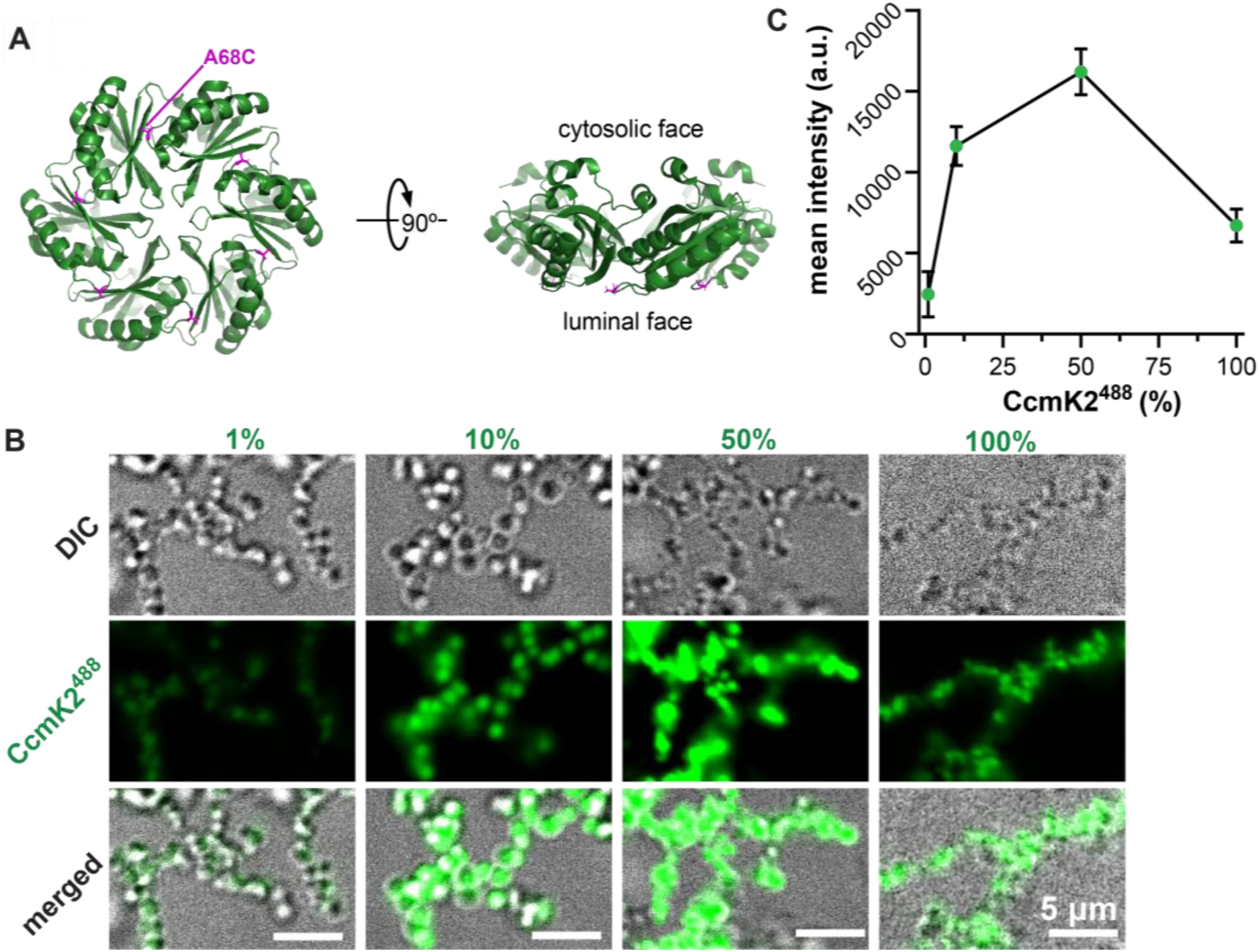
Particles assemble with increasing amounts of CcmK2^488^. **A.** CcmK2 structure showing the location of the A68C mutation (magenta) used for maleimide-dye conjugation. The cytosolic and luminal faces in native carboxysomes are labeled (PDB: 4OX7). **B.** DIC and fluorescence microscopy of particles assembled at the indicated percentage of CcmK2^488^. The remaining percentage is wildtype CcmK2 (total concentration = 40 µM). **C.** Mean fluorescence intensity of CcmK2^488^–decorated particles as a function of initial percentage of CcmK2^488^. Green dots and error bars indicate the mean ± standard deviation for n=3 technical replicates.

To determine whether direct conjugation of small molecules could also serve as an efficient method of loading significantly higher amounts of cargo, we imaged the assembly of CcmK2 particles mixed with 1, 10, 50, and 100% CcmK2^488^ (Figure 1B). CcmK2 particles assembled at every percentage of CcmK2^488^, but particle contrast in the differential interference contrast (DIC) channel and the fluorescence intensity greatly decreased at 100% CcmK2^488^, suggesting a negative impact on particle density when all CcmK2 is directly conjugated to Alexa Fluor 488. Still, the mean fluorescence intensity steadily increased from 1 to 50% CcmK2^488^ (Figure 1C). The data demonstrate that small molecules can be loaded via direct maleimide conjugation to CcmK2[A68C], but roughly half of the particle must be composed of wildtype CcmK2 for proper assembly and high loading capacity.

### CcmK2 particles non-specifically load small molecules (fluorescein) during assembly

The loading saturation and assembly inhibition observed with CcmK2^488^ likely arose because the mutated CcmK2^488^ subunits did not incorporate into particles as efficiently as wild-type CcmK2. We hypothesized that non-specific encapsulation of small molecules, rather than covalent fusion to a shell subunit, could enable higher loading capacities without compromising particle formation. Importantly, for certain biotechnological applications, cargo must be readily released from the particle after delivery; covalent attachment to shell proteins would prevent such release and limit functional utility. In this context, stochastic, non-covalent loading strategies offer a key advantage by enabling reversible cargo association and more flexible release compared to direct, irreversible conjugation approaches.

To test this hypothesis, we investigated whether CcmK2 particles could encapsulate the small fluorescent molecule fluorescein (MW = 376 Da; R_g_ = ∼ 0.5 nm). We developed a strategy in which cargo is loaded during particle assembly (Figure 2A, Methods). CcmK2 particles were assembled in samples containing increasing concentrations of fluorescein (Figure 2B-C). CcmK2^647^ was included in a 1:1000 ratio of CcmK2^647^:wildtype CcmK2 for visualization. We used microscopy and a fluorescein calibration curve (Figure S1A) to assess particle assembly and estimate the concentration and total number of molecules of fluorescein loaded into each particle.

**Figure 2.**
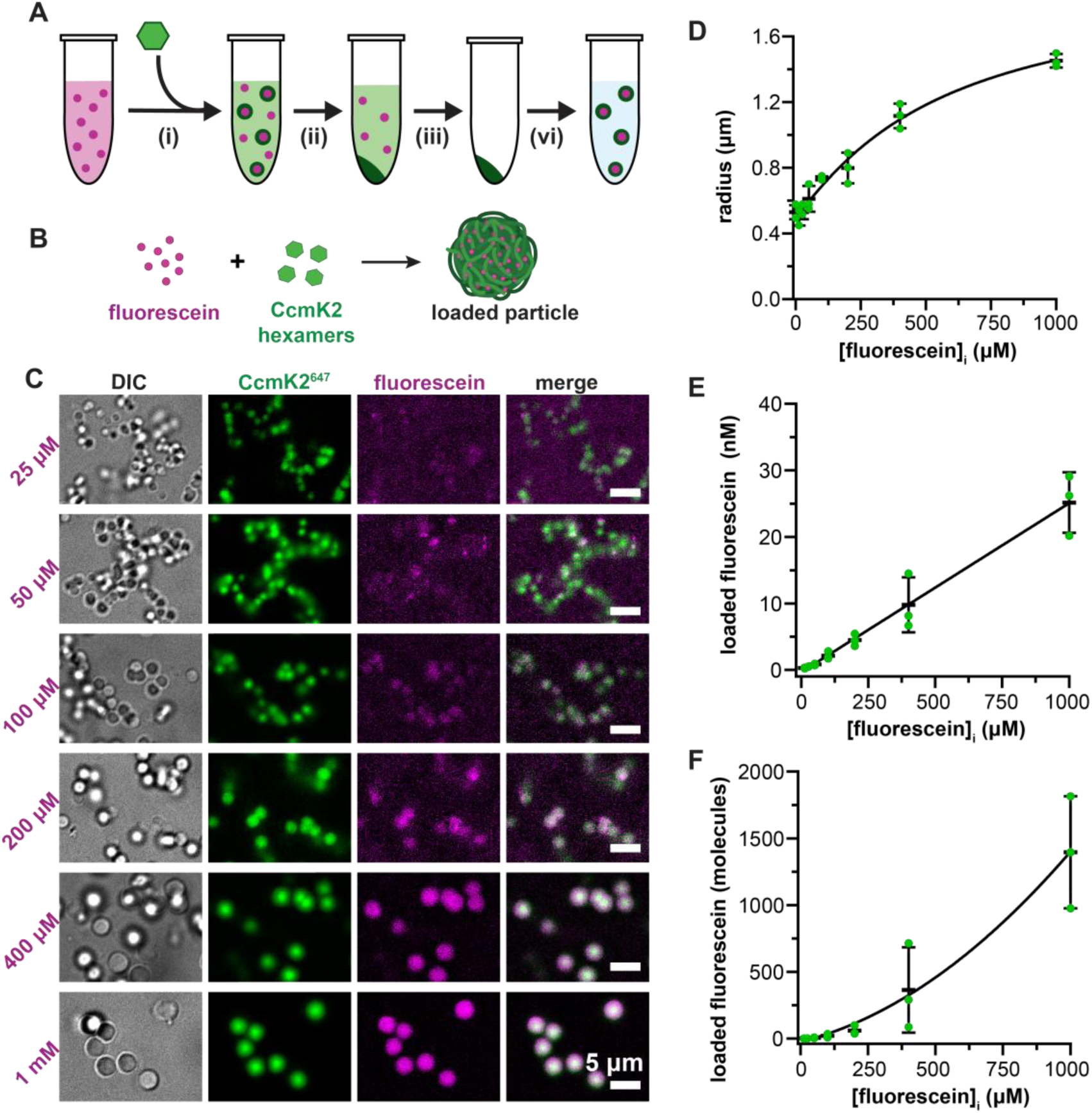
CcmK2 particles load fluorescein during assembly. **A.** Schematic of cargo loading during assembly: (i) soluble CcmK2 is added to assembly buffer containing cargo, (ii) loaded particles are pelleted via low-speed centrifugation. Free, unencapsulated cargo remains in the supernatant. (iii) The supernatant is removed and (iv) loaded particles are resuspended in cargo-free assembly buffer. **B.** Iconography of fluorescein loading during particle assembly. **C.** DIC and fluorescence microscopy images of particles assembled at the indicated initial concentration of fluorescein [fluorescein]_i_. **D-F.** Particle radius (D) concentration (E) and number (F) of fluorescein molecules per particle as a function of initial fluorescein concentration [fluorescein]_i_. Data were fit to a sigmoidal (D), linear (E), or quadratic (F) equation. Green dots indicate the means of individual replicates and black bars indicate mean ± standard deviation for n= 3 technical replicates.

Intriguingly, the size of CcmK2 particles increased with increasing initial concentrations of fluorescein (Figure 2D). But at a given fluorescein concentration, low polydispersity was maintained – that is, particles remained similarly sized. Particle diameter nearly tripled, rising from 1.0 ± 0.1 µm to 2.9 ± 0.1 µm, when particles were assembled in the presence of 12.5 µM-1 mM of fluorescein.

As the input concentration of fluorescein increased, both the mean fluorescence intensity (Figure S1B-C) and the concentration of loaded fluorescein (Figure 2E) increased linearly and did not reach saturation. The sum fluorescence intensity (Figure S1D-E) and the number of fluorescein molecules (Figure 2F) increased disproportionately with input concentration, rising from 1.3 ± 0.98 to 1400 ± 420 molecules per particle as the input concentration increased from 12.5 µM - 1 mM of fluorescein. The maximum loading capacity is likely higher as the number of fluorescein molecules did not plateau, suggesting that cargo saturation has not yet been reached. The nonlinear increase in total fluorescein suggests that loading follows a cooperative mechanism. This cooperativity likely arises from a positive feedback mechanism in which initial fluorescein incorporation promotes particle growth, which in turn increases the particle’s loading capacity and facilitates further cargo incorporation without altering packing density.

### CcmK2 particles non-specifically load protein cargo (mCherry) during assembly

After showing the ability of CcmK2 particles to load small fluorescent molecules both via direct conjugation and non-specifically during their assembly, we next investigated whether the particles could load significantly larger cargos, such as small proteins. We used the fluorescent protein mCherry (27 kDa; R_g_ = ∼ 2.2 nm) as a proof-of-concept - ∼ 70x larger in mass, ∼ 5x larger in R_g_, and ∼ 10x in volume compared to fluorescein. CcmK2 particles were assembled in the presence of increasing concentrations of mCherry (Figure 3A-B). CcmK2^488^ was included in a 1:1000 ratio of CcmK2^488^:CcmK2 for visualization. We again used microscopy and a calibration curve to assess particle assembly and to calculate mCherry loading from fluorescence intensities (Figure S2).

**Figure 3.**
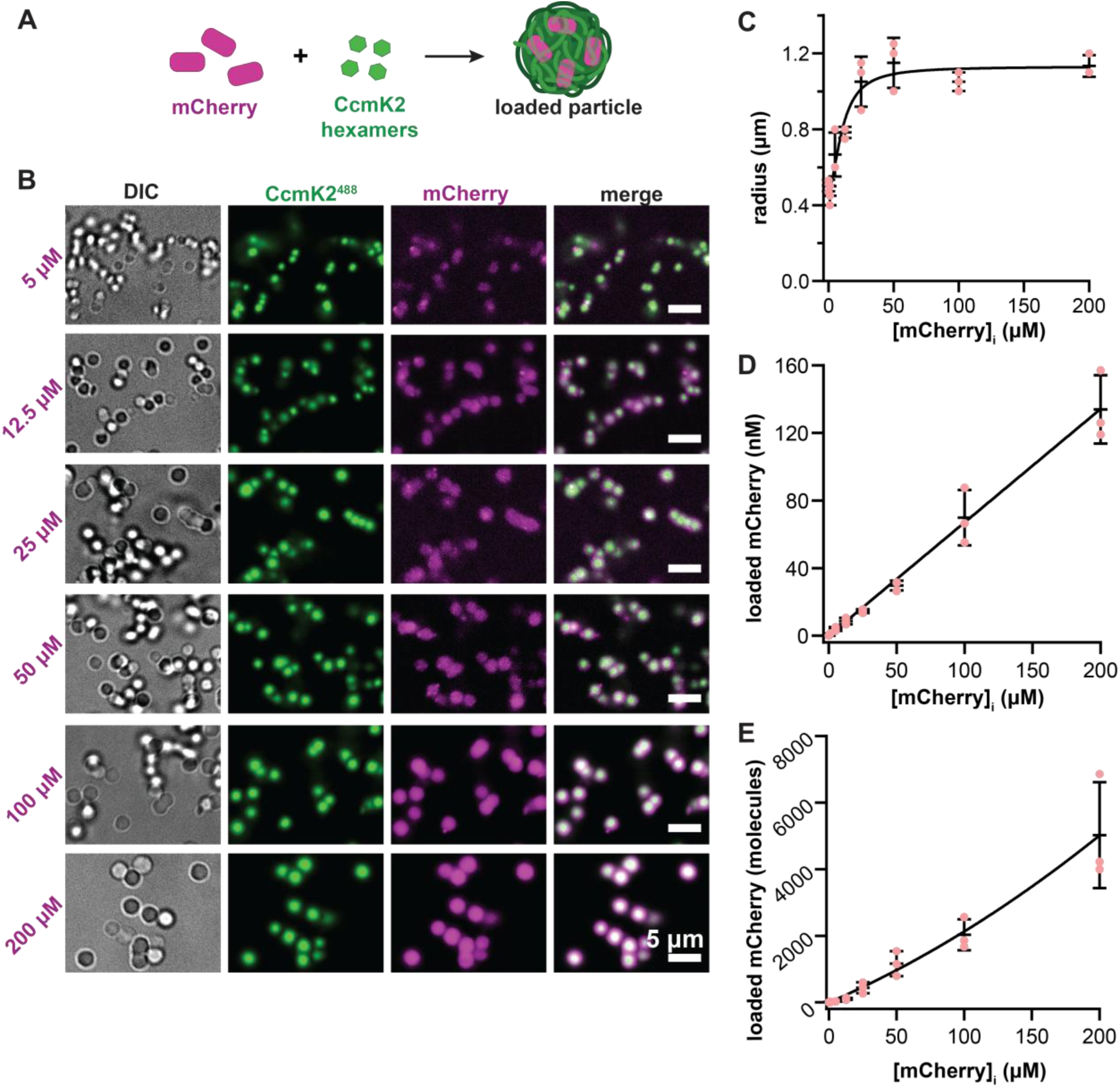
CcmK2 particles load mCherry during assembly. **A.** Iconography of mCherry loading during CcmK2 particle assembly. **B.** DIC and fluorescence microscopy images of particles assembled at the indicated initial concentration of mCherry. **C-E.** Concentration (C) and number (D) of mCherry molecules per particle, and particle radius (E) as a function of initial mCherry concentration [mCherry]_i_. Data were fit to a linear (C), quadratic (D), or sigmoidal (E) equation. Pink dots indicate the means of individual replicates and black bars indicate mean ± standard deviation for n= 3 technical replicates.

Across the mCherry titration, the size of CcmK2 particles once again increased; particles assembled to final diameters of 1.2 ± 0.5 µm at an initial concentration of 1 µM mCherry and plateaued at final diameters of 2.1 ± 0.2 µm with initial concentrations of 25 µM mCherry or greater (Figure 3C). The concentration of loaded mCherry increased linearly, rising from 1.23 ± 0.5 nM to 130 ± 20 nM (Figure 3D). As with fluorescein, the total number of loaded mCherry molecules increased non-linearly as a function of the initial mCherry concentration (Figure 3E). The non-linear increases in particle size and loaded molecules once again suggest cooperativity in the loading mechanism.

### Fusion of an encapsulation peptide to mCherry affects loading

To enhance cargo loading efficiency, we next turned to the native β-carboxysome assembly pathway. β-carboxysomes contain a scaffolding protein, CcmN, which consists of a C-terminal domain that interacts with core cargo proteins, a long intrinsically disordered linker, and a short N-terminal α-helix known as the encapsulation peptide (EP).^21^ The EP directly binds shell proteins such as CcmK2, thereby recruiting the shell to the carboxysome core. Importantly, studies across diverse BMCs have demonstrated that fusion of their EP to heterologous proteins is sufficient to drive encapsulation, establishing EPs as a portable targeting modules.^5,27–29^

We hypothesized that incorporating the EP could increase mCherry loading into CcmK2 particles by promoting specific interactions during assembly. To test this, we fused the EP to the C-terminus of mCherry via a flexible short GSGSGS linker, generating mCh-(GS)₃-EP, and assessed particle assembly and loading (Figure 4A). CcmK2 particles successfully assembled when incubated with 1–100 µM mCh-(GS)₃-EP (Figure 4B). Notably, particle abundance decreased progressively as the concentration of mCh-(GS)₃-EP increased. At 1 µM mCh-(GS)₃-EP, ∼ 200 particles were observed per representative field of view, whereas at 100 µM, only ∼10 particles were detected (Figure S3A). No particles were detected at 200 µM (Figure 4B).

**Figure 4.**
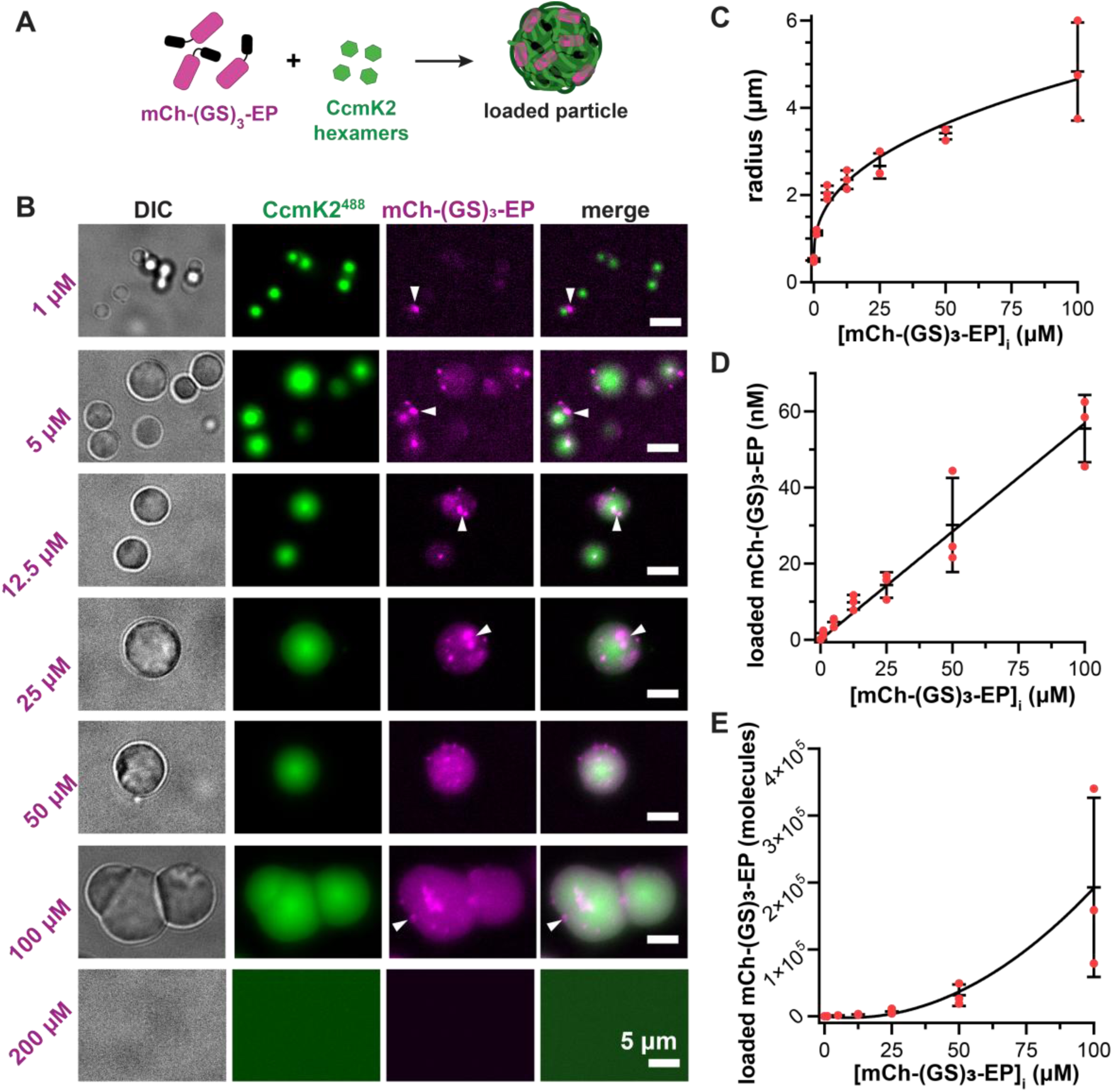
The encapsulation peptide affects mCherry-loading and particle assembly. Iconography of mCh-(GS)_3_-EP loading during CcmK2 particle assembly. **B.** DIC and fluorescence microscopy images of particles assembled at the indicated initial concentration of mCh-(GS)_3_-EP. White arrows indicate examples of mCh-(GS)_3_-EP puncta. Particles did not assemble in the presence of 200 µM mCh-(GS)_3_-EP. **C-E.** Particle radius (C) and concentration (D) and number (E) of mCh-(GS)_3_-EP molecules per particle as a function of initial mCh-(GS)_3_-EP concentration [mCh-(GS)_3_-EP]_i_. Data were fit to a sigmoidal (C), linear (D), or quadratic (E) equation. Red dots indicate means of individual technical replicates, and black bars indicate mean ± standard deviation for n = 3 technical replicates.

Strikingly, particle diameter quadrupled across the mCh-(GS)_3_-EP titration, rising from 1.2 ± 0.1 µm to 4.8 ± 1.1 µm (Figure 4B-C). While particle morphology was relatively uniform at low input concentrations of mCh-(GS)_3_-EP, size and morphology varied at high input concentrations. At 100 µM input, we observed unexpected morphologies such as hemispheres (Figure S3B), “wrinkly” uneven borders (Figure S3C), and fused particles (e.g., 100 µM mCh-(GS)_3_-EP in Figure 4B).

Unlike the uniform fluorescent signal from non-specifically loaded fluorescein and mCherry, there were several puncta and patches of brighter mCh-(GS)_3_-EP fluorescence colocalized within and on the surface of CcmK2 particles (Figure 4B, arrows). These patches and puncta are likely aggregated mCh-(GS)_3_-EP, which aligns with previous reports of fusion of the EP to heterologous proteins inducing aggregation.^21,30,31^

Despite the heterogeneous mCh-(GS)₃-EP fluorescence and accompanying changes in particle size distribution and morphology, mCh-(GS)₃-EP signal intensity remained positively correlated with its initial concentration (Figure S4). Across the range of 1–100 µM mCh-(GS)₃-EP, the concentration of mCh-(GS)₃-EP followed a linear trend (Figure 4D), while the number of loaded molecules increased disproportionately (Figure 4E). At 100 µM input, each particle contained nearly 3,000-fold more molecules of mCh-(GS)₃-EP than at 1 µM. Notably, this corresponded to ∼100-fold more molecules per particle than observed with 100 µM input of nonspecific mCherry. This non-linear increase in total cargo loading again indicates cooperativity, consistent with EP-mediated interactions enhancing recruitment during assembly. Moreover, the pronounced increase in particle size, altered morphology, and eventual assembly inhibition with increasing mCh-(GS)₃-EP concentration indicate that the EP also affects particle assembly.

### Linker length between mCherry and the EP modulates cargo loading

Although fusion of the EP increased cargo recruitment, the heterogeneous fluorescence and altered particle morphology observed with mCh-(GS)₃-EP suggested that geometric/steric constraints may limit optimal EP-CcmK2 interactions. The EP of *S. elongatus* CcmN is separated from its cargo-binding domain by a 21-amino-acid intrinsically disordered linker^21^, likely providing sufficient flexibility to enable both productive cargo incorporation and shell assembly. We therefore hypothesized that the relatively short linker in mCh-(GS)₃-EP may sterically hinder EP accessibility, perturbing assembly and promoting uneven cargo distribution.

To test whether increasing flexibility would improve loading behavior, we doubled the linker length to generate mCh-(GS)₆-EP and used particle-associated fluorescence intensity as a readout of cargo loading (Figure 5A, S5). CcmK2 particles assembled in the presence of 1–100 µM mCh-(GS)₆-EP, but assembly was once again inhibited at 200 µM (Figure 5B). Across this titration range, particle size and morphology were markedly more uniform than those observed with mCh-(GS)₃-EP (Figure 5C). Particle diameter increased with increasing initial mCh-(GS)₆-EP concentration but plateaued above 25 µM, suggesting a transition to cargo-saturated particle assemblies.

**Figure 5.**
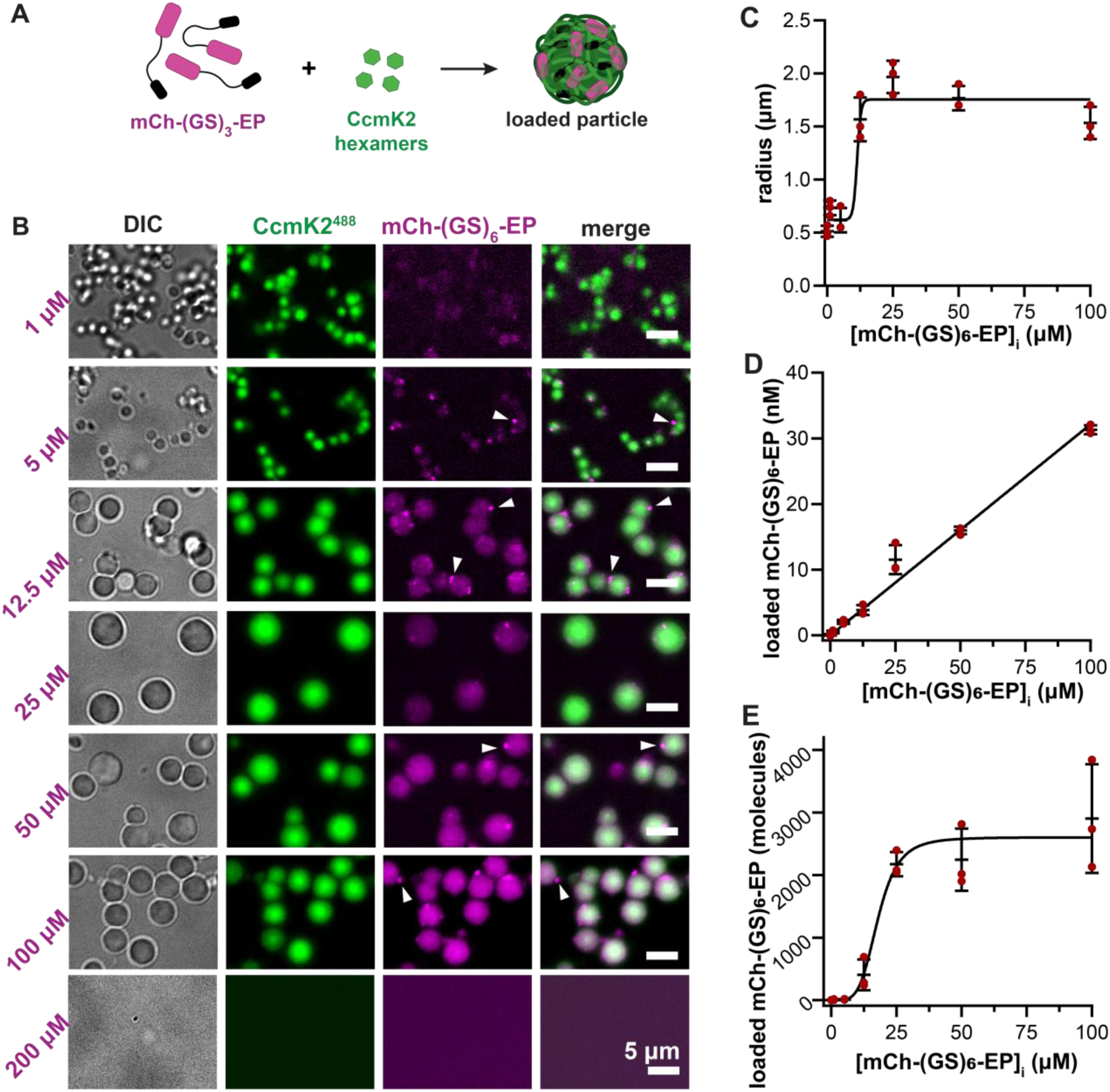
The linker length between mCherry and the encapsulation peptide modulates loading. **A.** Iconography of mCh-(GS)_6_-EP loading during CcmK2 particle assembly. **B.** DIC and fluorescence microscopy images of particles assembled at the indicated initial concentration of mCh-(GS)_6_-EP. White arrows indicate examples of mCh-(GS)_6_-EP puncta. Particles did not assemble in the presence of 200 µM mCh-(GS)_6_-EP. **C-E.** Particle radius (C) and concentration (D) and number (E) of mCh-(GS)_6_-EP molecules per particle as a function of initial mCh-(GS)_6_-EP concentration [mCh-(GS)_6_-EP]_i_. Data were fit to a sigmoidal (C,E) or linear (D) equation. Dark red dots indicate means of individual technical replicates, and black bars indicate mean ± standard deviation for n = 3 technical replicates.

Fluorescence from mCh-(GS)₆-EP was also more evenly distributed, with occasional puncta within particles and at their surface (Figure 5B, arrows). As the input concentration of mCh-(GS)₆-EP increased, the concentration of mCh-(GS)₆-EP increased linearly (Figure 5D). The number of molecules per particle followed a sigmoidal trend, rising from 8 ± 5 to 2900 ± 870 as input concentration increased from 1 to 100 µM (Figure 5E).

Because linker length is the only variable distinguishing mCh-(GS)₃-EP and mCh-(GS)₆-EP, these results demonstrate that linker length is a key determinant of particle assembly dynamics, cargo distribution, loading density, and maximal loading capacity. Modulating linker length therefore provides a tunable parameter for engineering cargo encapsulation within CcmK2 particles.

### The SpyTag-SpyCatcher system immobilizes cargo onto CcmK2 particles

Several studies have demonstrated that while a BMC’s native EP is sufficient for recruiting heterologous cargo, other anchoring systems, like the SpyTag/SpyCatcher technology,^22^ achieve more efficient cargo loading.^13,14,29,32^ Recently, by engineering an internal SpyCatcher (SC) fusion within the hexameric shell protein CsoS1A of the α-carboxysome, the Liu lab successfully recruited SpyTagged (ST) cargo to recombinant α-carboxysomes.^29^ We therefore engineered an internal SC fusion into the heterologous site of CcmK2 to form K2^SC^ (Figure 6A). In parallel, we constructed mCherry with an N-terminal His-ST fusion (ST-mCh; MW= 31 kDa) to function as a directed cargo for encapsulation and immobilization within CcmK2 particles.

**Figure 6.**
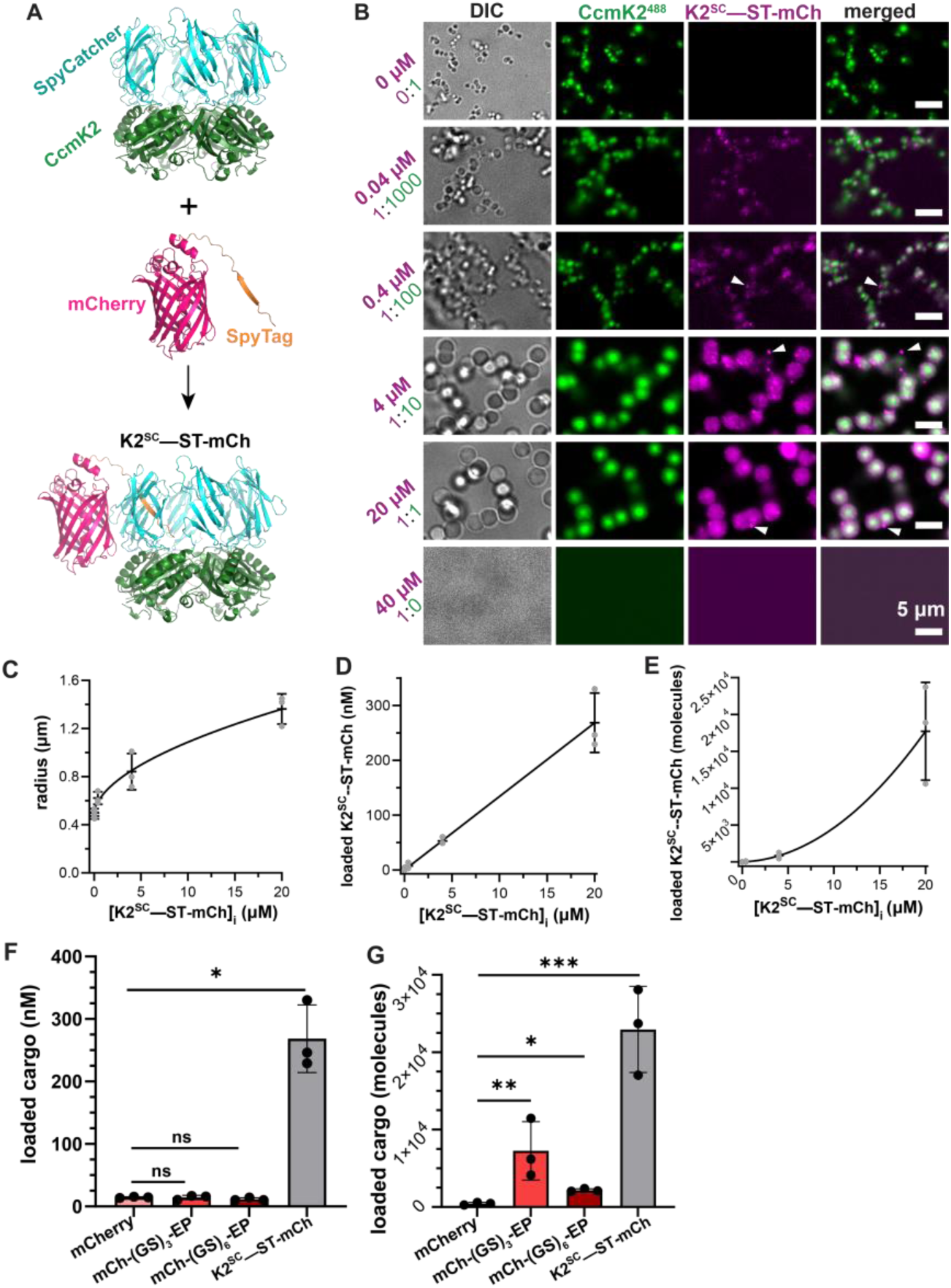
The SpyCatcher/SpyTag system enables protein loading via direct conjugation. **A.** SpyCatcher (cyan) was fused internally to each subunit of the CcmK2 hexamer (green) to general CcmK2^SC^. SpyTag (orange) was fused to the N-terminus of mCherry (hot pink) to generate ST-mCh. ST-mCh associates with CcmK2^SC^. Five other ST-mCh molecules (not shown) associate to form the full K2^SC^—ST-mCh complex. Models were generated using AlphaFold3.^39^ **B.** DIC and fluorescence microscopy images of CcmK2 particles assembled at the indicated concentration of K2^SC^—ST-mCh. The molar ratio of K2^SC^—ST-mCh:CcmK2 is also indicated (total concentration = 40 µM). **C-E.** Particle radius (C) and concentration (D) and number (E) of K2^SC^—ST-mCh molecules per particle as a function of initial K2^SC^—ST-mCh concentration [K2^SC^—ST-mCh]_i_. Data were fit to a sigmoidal (C), linear (D), or quadratic (E) equation. Gray dots indicate means of individual replicates, and black bars indicate mean ± standard deviation (SD) for n = 3 technical replicates. **F.** Concentrations of loaded mCherry variants at initial concentrations of 25 µM (mCherry, mCh-(GS)_3_-EP, mCh-(GS)_6_-EP) or 20 µM (K2^SC^—ST-mCh). Dots indicate means of individual replicates, and bars and error bars indicate mean ± SD for n=3 technical replicates. Statistical analyses were performed using unpaired two-tailed t-tests with Welch’s correction.

Prior to investigating cargo loading, we confirmed that K2^SC^ and ST-mCh could form a complex (hereafter, K2^SC^—ST-mCh) (Figure S6, Methods). We tested the assembly of particles with increasing concentrations of K2^SC^—ST-mCh by adjusting the ratio of wild-type CcmK2:K2^SC^—ST-mCh, keeping the total concentration at 40 µM. Despite the significant addition of the SC domain (9 kDa) to each CcmK2 monomer (11 kDa), particles still faithfully assembled in the presence of 0.1 to 50% K2^SC^—ST-mCh, but not at 100% K2^SC^—ST-mCh (Figure 6B). The final diameters of the CcmK2 particles were similar at a specific K2^SC^—ST-mCh concentration, and nearly tripled (1.0 ± 0.2 to 2.7 ± 0.4 µm) across the titration (Figure 6C).

Across the titration of K2^SC^—ST-mCh, fluorescence intensity increased (Figure S7) and was evenly distributed with rare puncta (Figure 6B, arrows); notably fewer than those seen with mCh-EP proteins. As the input concentration of K2^SC^—ST-mCh increased, the calculated concentration increased linearly, reaching 270 ± 50 nM at 20 µM input (Figure 6D). The calculated number of encapsulated K2^SC^—ST-mCh molecules increased non-linearly with input concentration, rising from 15 ± 9 to 23,000 ± 5,600 molecules per particle (Figure 6E). These results again suggest cooperative loading behavior, which is consistent with the design of this construct, as direct conjugation of CcmK2 to mCherry intrinsically links cargo incorporation to particle assembly.

Relative to the initial cargo concentration, K2^SC^—ST-mCh facilitated significantly more cargo loading than the other mCherry variants, both in terms of concentration (∼two to nine-fold higher than other variants) (Figure 6F) and the number of molecules per particle (∼ 3 to 52-fold more than other variants) (Figure 6G). These data demonstrate that the ST/SC system efficiently loads immobilized cargo onto CcmK2 particles.

### Preassembled CcmK2 particles are permeable to small dyes and proteins

After assessing cargo loading during CcmK2 particle assembly, we determined how well cargo could permeate into preassembled particles (see Methods). First, we tested small molecule permeation using 200 µM of fluorescein (Figure 7A). Notably, particle diameter did not change (Figure 7B). Over time, the concentration of particle-associated fluorescein and the number of molecules loaded per particle increased but plateaued above 20 h (Figure 7C-D). By 120 h, the concentration of permeated fluorescein (10.5 ± 1.7 µM) slightly exceeded the concentration that was loaded during assembly at 200 µM input (4.5 ± 0.9 µM) (Figure 7E). Total fluorescein permeation, however, did not exceed the number of molecules loaded during assembly at 200 µM input (Figure 7F).

**Figure 7.**
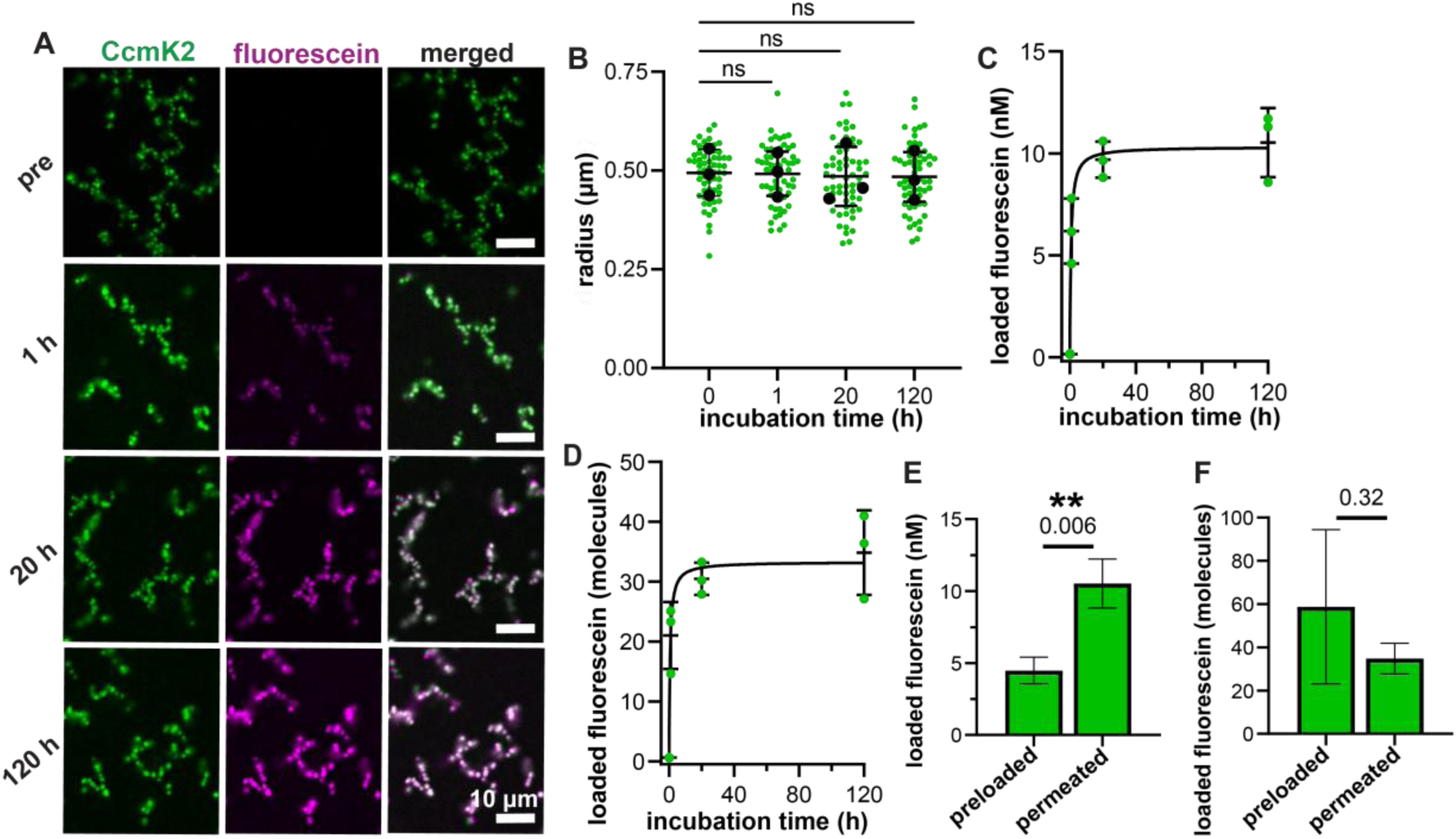
Fluorescein permeates into pre-assembled CcmK2 particles. **A.** Fluorescence microscopy images of fluorescein molecules permeated into pre-assembled CcmK2 particles at the indicated incubation time. “pre” indicates particles before fluorescein was added. **B.** Particle diameter after incubation with fluorescein. Green dots represent individual data points across 3 replicates. Black dots indicate means of each replicate, and black bars represent the overall mean ± standard deviation (SD) for n = 3 technical replicates. Statistical analyses were performed using one-way ANOVA. **C-D.** Concentration **(C)** and number **(D)** of fluorescein molecules per particle as a function of incubation time. Green dots indicate the means of individual replicates and black bars indicate total mean ± SD for n=3 technical replicates. Data were fit to a hyperbola. **E, F.** Concentration (**E**) and number **(F)** of fluorescein molecules per particle when 200 µM fluorescein was provided during (preloaded) or after (permeated) particle assembly. Data represent mean ± SD for n = 3 technical replicates. Statistical analyses were performed using unpaired two-tailed t-tests with Welch’s correction.

We also tested how much mCherry, mCh-(GS)_3_-EP, and mCh-(GS)_6_-EP, could permeate into CcmK2 particles by adding 100 µM of each cargo to preassembled particles (Figure 8A). Once again, the size of preassembled particles did not change over time (Figure 8B). From 1 to 24 h, the particle-associated concentration and the number of molecules loaded per particle increased for all mCherry cargos without plateauing (Figure 8C-D). By 24 h, the concentration of all permeated mCherry variants equaled or exceeded the concentration of molecules loaded during assembly when 100 µM cargo was input (Figure 8E). However, total permeation was one to three orders of magnitude lower than the number of molecules loaded during assembly (Figure 8F).

**Figure 8.**
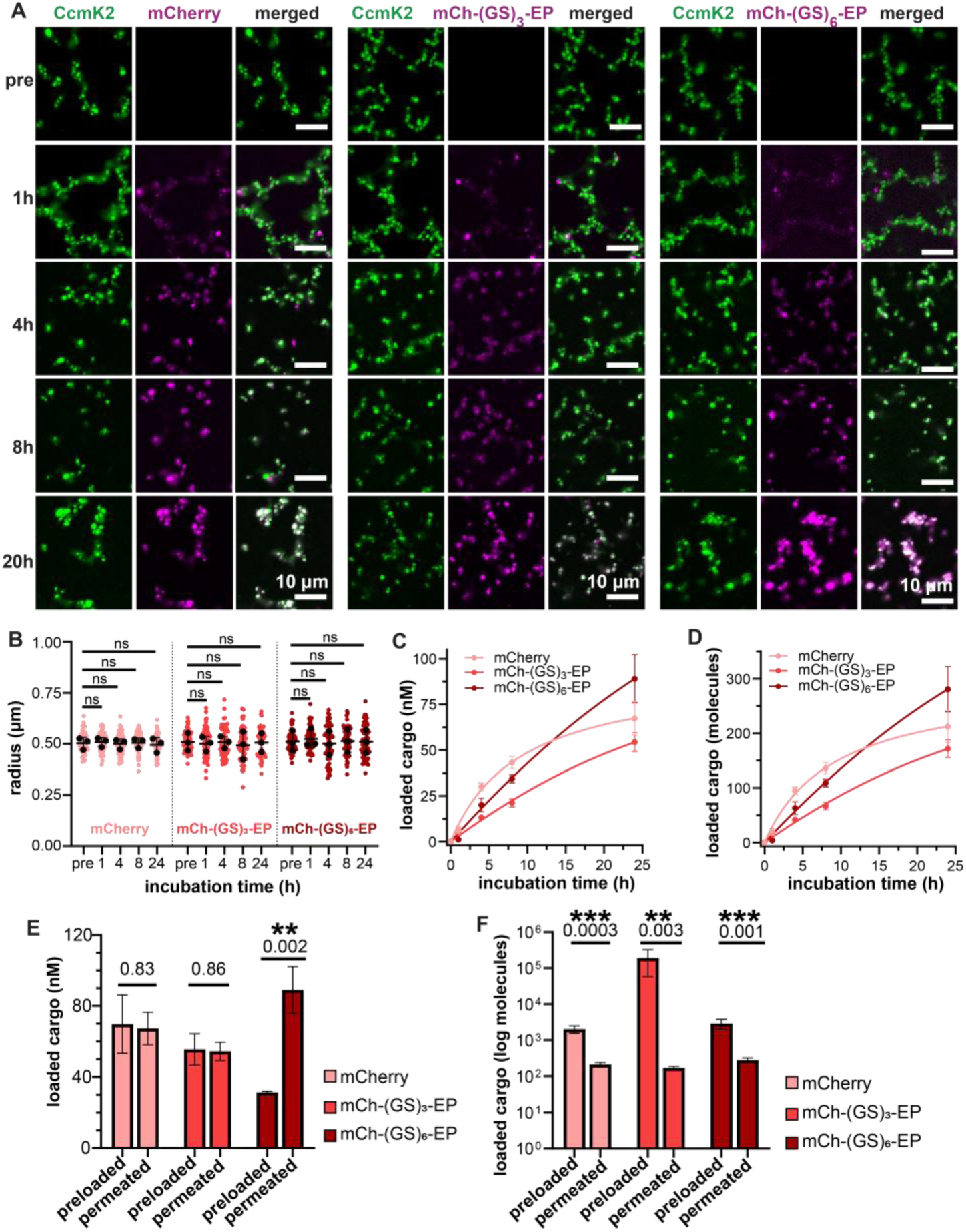
mCherry variants permeate into pre-assembled CcmK2 particles. **A.** Fluorescence microscopy images of mCherry (left), mCh-(GS)_3_-EP (middle), and mCh-(GS)_6_-EP (right) permeated into pre-assembled CcmK2 particles at the indicated incubation time. “pre” indicates particles before cargo was added. **B.** Particle diameter after incubation with the indicated cargo. Colored dots represent individual data points across 3 replicates, black dots indicate means of each replicate, and black bars represent total mean ± standard deviation (SD); n=3 technical replicates. Statistical analyses were performed using one-way ANOVA. **C-D.** Concentration **(C)** and number **(D)** of cargo molecules per particle as a function of incubation time. Data represent mean ± SD for n=3 technical replicates. mCherry was fit to a hyperbola and mCh-EP variants were fit to quadratic equations. **E, F.** Concentration (**E**) and number **(F)** of cargo molecules per particle when 100 µM cargo was provided during (preloaded) or after (permeated) particle assembly. Data represent mean ± SD for n = 3 technical replicates. Statistical analyses were performed on raw (E) or log-transformed (F) data using unpaired two-tailed t-tests with Welch’s correction.

Together, we draw the following conclusions: (1) CcmK2 particles are permeable to both small molecules (fluorescein-380 kDa) and small proteins (mCherry-27 kDa), (2) permeation of cargo into pre-assembled CcmK2 particles does not affect their size, (3) targeting modules do not greatly affect cargo permeation into CcmK2 particles, and (4) cargo permeation is likely driven by a stochastic mechanism, such as diffusion-limited penetration into particles, while cargo loading during assembly is governed by a cooperative mechanism.

## DISCUSSION

In this report, we demonstrate that CcmK2 particles can encapsulate diverse cargo, including small molecules, small proteins, and large protein complexes, via direct conjugation and passive or targeted loading strategies. Fusion of mCherry to the native carboxysome EP enhances particle loading, highlighting the utility of native targeting modules for directing cargo to BMC-based platforms. Among the strategies tested, immobilization of mCherry to CcmK2 using the ST/SC system yields the highest cargo-loading efficiency. We further identified the length of the linker connecting cargo to the EP, as well as the order of cargo introduction, to be tuners of final particle size. Collectively, these results establish a framework for controlling cargo loading and accessibility, as well as particle properties, which supports the development of CcmK2 particles as customizable platforms for biotechnological applications.

### Cooperative cargo loading is driven by nonspecific and multivalent interactions

We developed two general strategies for loading cargo into CcmK2 particles: simultaneous cargo loading during particle assembly (Figure 9A) and cargo permeation into preassembled particles (Figure 9B). With equivalent input concentrations of cargo, simultaneous loading during assembly loads one to three orders of magnitude more molecules of cargo per particle than permeation. Additionally, increasing cargo input during assembly leads to larger particles, whereas particle size remains unchanged during permeation. Together, these observations indicate that cargo loading during assembly is a cooperative process that actively promotes particle growth, enabling substantially greater loading than diffusion-driven permeation into preassembled particles.

**Figure 9.**
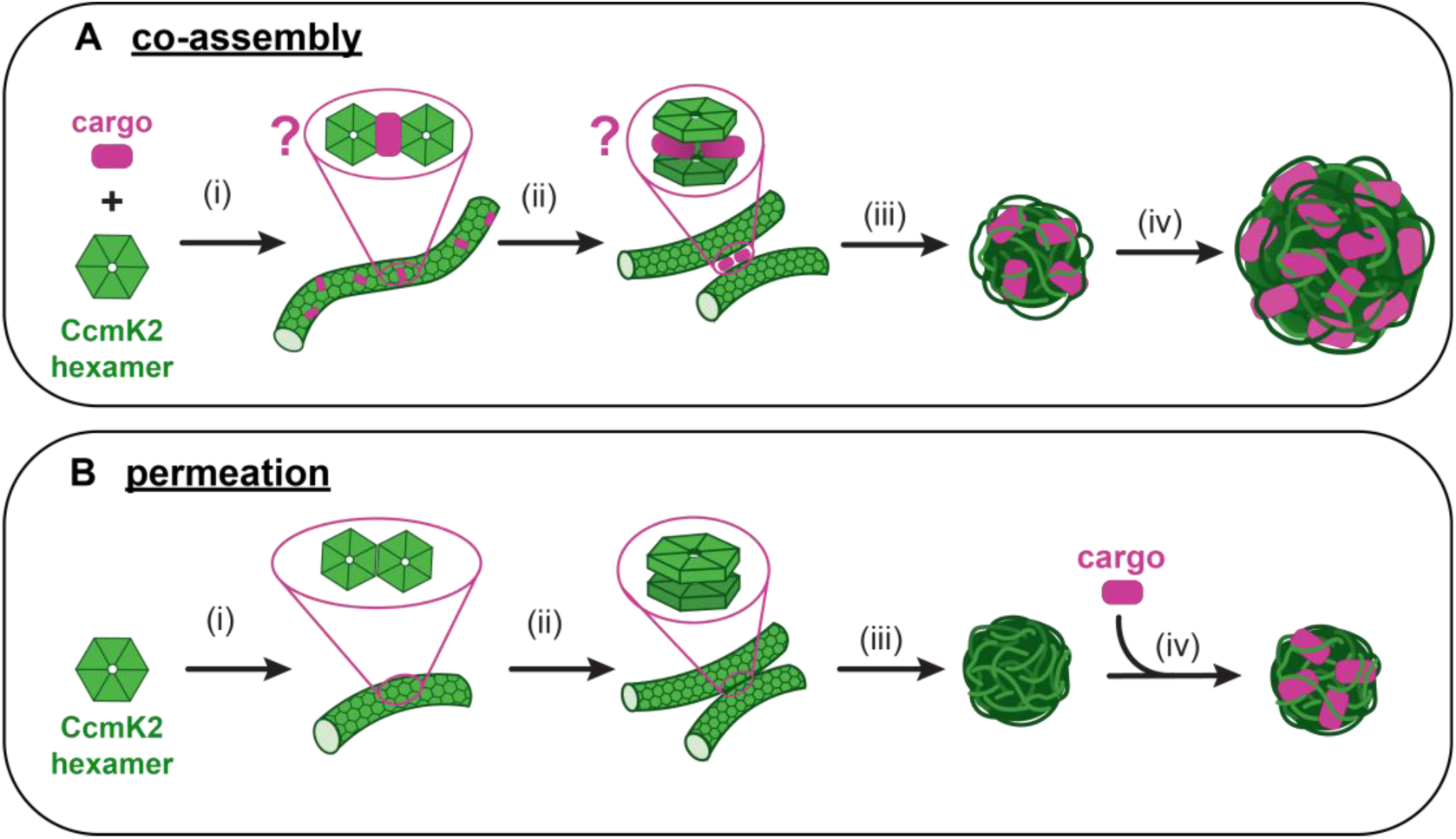
Proposed mechanisms for cargo loading. **A.** When cargo is introduced during assembly, (i) cargo may alter CcmK2 hexamer side-side interactions thus affecting nanotube assembly. (ii) Cargo may also affect nanotube clustering by altering CcmK2 face-face interactions. (iii) The upstream effects of cargo modify the assembling nanotube meshwork, (iv) facilitating the formation of larger particles and increased cargo loading. **B.** In the absence of cargo, particles assemble as previously reported.^17^ (i) CcmK2 hexamers tile into nanotubes via side–side hexamer contacts. (ii) CcmK2 nanotubes cluster by interacting via face–face hexamer contacts. (iii) CcmK2 nanotubes continue to cluster radially to form the final mesh-like particle. (iv) Cargo added to preassembled particles loads via permeation without affecting particle size or architecture.

We propose that this cooperativity arises from a co-assembly mechanism in which weak, non-specific interactions between cargo and assembling CcmK2 structures generate a partitioning-like effect that positively feeds back on particle assembly and cargo incorporation (Figure 9A). Consistent with this model, chemically distinct cargos (mCherry and fluorescein) exhibit similar loading behavior: total cargo loading increases nonlinearly with input concentration, while loaded cargo concentration scales linearly. These observations disagree with strong, saturable binding and instead support a framework in which multivalent, low-affinity interactions (e.g., electrostatics, hydrophobic effects, and molecular crowding) collectively promote co-assembly of CcmK2 particles and cargo.

The effects of the EP further support this model, while also implicating a role for more specific interactions. Fusion of the EP to mCherry increases total cargo loading without significantly altering the loaded cargo concentration, indicating enhanced co-assembly rather than increased packing density. The amphipathic nature of the EP likely enables it to provide additional non-specific, multivalent interactions (i.e., hydrophobic and electrostatic) while specific EP-CcmK2 interactions may stabilize cargo incorporation.

Importantly, this cooperative effect must be reflected in the proposed hierarchical assembly of CcmK2 particles, but the specific mechanism remains unresolved. Increased cargo loading could be accommodated through multiple, non-mutually exclusive effects. Cargo may affect nanotube assembly by altering the CcmK2 interhexamer tiling interface, thereby changing nanotube curvature, flexibility, or elongation dynamics (Figure 9A(*i*)). Another possibility is that cargo may influence higher-order particle organization, where steric effects and/or multivalent interactions between cargo and nanotubes reorganize, disrupt, or enhance nanotube-nanotube contacts, expanding the meshwork and promoting the formation of larger particles (Figure 9A(*ii*)). In all cases, the key feature is that cargo packing density remains constant, while particle architecture adjusts to increase particle size and accommodate greater total loading. Defining the assembly steps that are sensitive to cargo will be essential for establishing the mechanistic basis of cooperative loading and will inform the rational engineering of CcmK2 particles with tunable size, architecture, and cargo capacity.

### The EP linker length modulates valency during assembly

We also compared the efficacy of the EP in directing cargo to CcmK2 particles when fused to mCherry via a short (mCh-(GS)_3_-EP) or a long (mCh-(GS)_6_-EP) linker. mCh-(GS)_3_-EP loads up to 200,000 ± 130,000 molecules per particle without saturating but severely alters particle morphology. mCh-(GS)_6_-EP rescues canonical particle morphology but saturates loading capacity around 2,900 ± 900 molecules per particle. Interestingly, at high enough concentrations of input cargo, both (GS)_3_ and (GS)_6_ EP fusions prevented particle assembly, suggesting that the EP itself destabilizes CcmK2 self-interactions involved in typical particle assembly. The contrasting loading results, however, demonstrate that the length of the linker between the EP and its cargo modulates the co-assembly of CcmK2 and cargo.

We propose that because both mCherry and the EP evidently provide multivalency during particle assembly, the decreased flexibility of the (GS)_3_ linker creates sites of highly localized avidity that increase the effective binding strength between CcmK2 and the cargo, stabilizing both normal and aberrant assemblies. Moreover, it is unlikely that the linkers themselves provide multivalency as they are composed only of flexible glycine–serine repeats lacking defined interaction motifs. Thus, doubling the linker length likely decreases local avidity, weakens the networking interactions that drive assembly, and increases the excluded volume. These effects are consistent with the lower cargo loading capacity and saturation behavior of mCh-(GS)_6_-EP when sequestered by CcmK2 particles. The linker between the EP and cargo is therefore a key engineering target for modulating particle assembly and loading capacity.

### EP-mediated loading provides insight into the native encapsulation mechanisms of β-carboxysomes

The implications of this study extend beyond engineered cargo loading to native β-carboxysome assembly. Our results suggest that EP-shell interactions are sufficient for cargo recruitment but must be balanced to support productive assembly. Increasing concentrations of mCh-(GS)₃-EP reduced particle abundance, and high concentrations of either EP fusion inhibited assembly, indicating that excess or mislocalized EP interactions disrupt shell formation. These observations support a model in which cargo encapsulation emerges from a balance of shell-shell and shell-EP interactions, opposed to EP affinity alone.

This interpretation is consistent β-carboxysome stoichiometry, in which each organelle contains only ∼8 copies of CcmN (the EP-containing scaffold protein), despite the presence of ∼2,000 shell oligomers. This disparity suggests that CcmN is not a major structural contributor to the shell but instead transiently nucleates or guides encapsulation.^33^ Our findings extend this view by suggesting that EP abundance is not merely minimized because additional copies are unnecessary, but because excessive EP engagement may actively disrupt assembly, effectively “poisoning” productive assembly pathways.

A key distinction between our system and native assembly further supports this model. *In vivo*, core proteins, including CcmN, first form a condensed pro-carboxysome that concentrates EPs and promotes coordinated shell recruitment.^21,34^ In contrast, EPs in our system are distributed on soluble cargo, allowing dispersed and uncoordinated EP–CcmK2 interactions that may influence CcmK2-CcmK2 interactions without promoting higher-order structure. Under these conditions, elevated EP levels may sequester shell proteins into nonproductive complexes. This distinction highlights the importance of spatial organization and phase behavior in regulating EP function during native carboxysome biogenesis.

### Developing CcmK2 particles as molecular scaffolds for biocatalysis

Here we used fluorescent molecules as proof-of-concept cargo to develop a suite of encapsulation strategies for CcmK2 particles across multiple biotechnologically relevant scales. While these cargos established the feasibility of cargo encapsulation, an important next step is to demonstrate whether these particles can function as scaffolds for enzyme immobilization and biocatalysis; likely in a cell free context. Recent studies have demonstrated that immobilizing enzymes to BMC-based scaffolds improves catalytic performance and stabilizes enzymes.^14,15,35,36^ However, few systems offer the combination of controlled reversible assembly, tunable size, and modular cargo loading - features that are highly desirable for broad biotechnological applications. Future work should therefore focus on whether enzymes loaded into CcmK2 particles retain optimal catalytic activity and benefit from loading. Once feasibility is established, the cargo-loading strategies we describe here may enable the co-immobilization of multiple enzymes for the modular spatial organization of multi-step reactions or substate channeling. These efforts would extend our current proof-of-concept findings toward the development of CcmK2 particles as versatile molecular scaffolds for biocatalysis.

## Materials and methods

### Protein Expression and Purification

Wildtype and mutant variants of CcmK2 from *S. elongatus* (PCC 7942) were purified as previously described.^17^ mCherry variants fused to an N-terminal His-tag were recombinantly expressed in *E. coli* BL21-AE (Invitrogen) from a pET11b vector. Cells were grown in LB + carbenicillin (100 μg/mL). One-liter cultures were inoculated using overnight cultures at a 1:100 dilution, grown to an OD600 of 0.6–0.8 and cooled on ice prior to expression induction using 1 mM isopropyl ß-D-1-thiogalactopyranoside (IPTG). Cultures were grown at 16 °C for 20 h, pelleted, flash-frozen in liquid nitrogen and stored at – 80 °C.

The following buffers were used to purify all mCherry variants: Buffer A (20 mM Tris-HCl [pH 7.8], 150 mM KCl, 10% glycerol, 20 mM imidazole, 5 mM BME), Lysis Buffer (buffer A + 0.05 mg/mL lysozyme, protease inhibitor, 5 mM BME), High Salt Buffer (20 mM Tris-HCl [pH 7.8], 1 M KCl, 10% glycerol, 20 mM imidazole), Buffer B (20 mM Tris-HCl [pH 7.8], 150 mM KCl, 10% glycerol, 1 M imidazole, 5 mM BME), and Storage Buffer (20 mM Tris-HCl [pH 7.8], 50 mM KCl, 10% glycerol, 2 mM DTT). Cell pellets were resuspended in 40 mL Lysis Buffer, and sonicated with cycles of 10 s on, 20 s off at 50% power for 6 min. The lysates were centrifuged at 19,000 rcf at 4 °C for 30 min.

Clarified lysates were decanted, syringe filtered (0.45 µm), and loaded onto a 5 mL HP His-TRAP (Cytiva) column equilibrated in Buffer A. The column was washed with five column volumes of High Salt Buffer, 5 column volumes of Buffer A, and 5 column volumes of 5% Buffer B. Elution was performed using a 5–100% gradient of Buffer B via an AKTA Pure system (Cytiva). Peak fractions were pooled, syringe filtered (0.2 μm) and passed over a size exclusion column (HiLoad 16/600 Superdex 200 pg; Cytiva) equilibrated in Storage Buffer. Peak fractions were pooled and concentrated to 5-10 mg/mL. Aliquots were flash-frozen and stored at −80 °C.

### CcmK2 Particle Assembly

CcmK2 (in Storage Buffer) was diluted into the assembly buffer (20 mM KPi [pH 6.2] and 150 mM KCl) to a final concentration of 40 μM unless otherwise stated. For fluorescent particle assembly, Alexa Fluor 488- or 647-labeled CcmK2[A68C] was mixed in a 1:1000 molar ratio of labeled- to dark-CcmK2 prior to dilution into the assembly buffer. Reactions were incubated at room temperature for 60 min prior to further experimentation.

### Design and Assembly of the CcmK2-SpyCatcher/SpyTag-mCherry complex

CcmK2^SC^ was generated by inserting ΔN1ΔC2SpyCatcher003^37^ flanked by GSGGS linkers into the flexible loop on the concave surface of CcmK2, between second α-helix and the fourth β-sheet (between residues 69 and 70). ST-mCherry was generated by fusing SpyTag003^23^ with an N-terminal hexahistidine tag to the N-terminus of mCherry with a GSGESG linker.

CcmK2^SC^ (20 µM) was incubated with two-fold molar excess of ST-mCherry (40 µM) in reaction buffer (20 mM potassium phosphate [pH 7.8] and 150 mM KCl) at room temperature for 5 minutes. The reaction was quenched by addition of ice-cold buffer followed by immediate spin filtration using a 100 kDa molecular weight cut-off Amicon Ultra-0.5 centrifugal filter device. Excess unreacted ST-mCherry was removed via five to six successive rounds of spin filtration and additions of reaction buffer to the concentrate. The CcmK2^SC^—ST-mCherry complex was concentrated to ∼ 200 µM and immediately used for cargo loading experiments.

### Loading cargo into CcmK2 particles

For the loading of fluorescein, mCherry, mCh-(GS)_3_-EP, and mCh-(GS)_6_-EP during particle assembly, cargo was included in the assembly buffer at the indicated concentration. CcmK2 was then diluted into the assembly buffer and reactions were incubated at room temperature for 60 minutes. Samples were spun at 16,000 rcf for 10 min, the supernatant was removed, and particles were resuspended in cargo-free assembly buffer. This washing process was performed three times to remove all free, unencapsulated cargo.

For loading CcmK2^488^ and CcmK2^SC^—ST-mCh, CcmK2^488^ or CcmK2^SC^—ST-mCh were premixed with wildtype CcmK2 in storage buffer at the indicated molar ratios. The mixtures of wildtype and variant CcmK2 were diluted into assembly buffer (40 µM final) and reactions were incubated at room temperature for 60 minutes and washed three times.

For cargo-loading after particle assembly, particles were spun at 16,000 rcf at room temperature for 10 min, the supernatant was removed, and particles were resuspended in assembly buffer plus cargo at the indicated concentration. Samples were incubated at room temperature for the indicated time and washed three times.

### Differential Interference Contrast (DIC) and Fluorescence Microscopy

Samples were transferred to 16 well CultureWells (Grace Biolabs) blocked with 5% Pluronic F-127. Imaging was performed using a Nikon Ti2-E motorized inverted microscope (100 × DIC objective and DIC analyzer cube) controlled by NIS Elements software with a Transmitted LED Lamp house and a Photometrics Prime 95B Back-illuminated sCMOS Camera. Alexa Fluor 488 and fluorescein sodium signals were acquired using a ‘GFP’ filter set [excitation/emission 488/519]. Alexa Fluor 647 signal was acquired using a ‘Cy5’ filter set [excitation/emission 640/670]. mCherry signal was acquired using a ‘TexasRed’ filter set [excitation/ emission 560/630].

Image analysis was performed using Fiji v 1.54f. Diameters of CcmK2 particles were analyzed by manually drawing line measurements across the diameters of particles in the DIC channel. Twenty measurements for were taken for each replicate. Mean and sum fluorescence intensities of particles were acquired using the TrackMate plugin in Fiji.^38^ Data were exported, further tabulated, graphed, and analyzed using GraphPad Prism 10.4.1 for Windows (GraphPad Software, Boston, Massachusetts USA, www.graphpad.com).

### Generation of fluorescein and mCherry calibration curves

Fluorescein sodium (0-1000 µM) or His-mCherry (0-1000 µM) in 20 µM potassium phosphate [pH 6.2] and 150 mM KCl was imaged using the ‘GFP’ or ‘TexasRed’ filter set, respectively (see section 1.6.5 above). For each of three replicates, 20 µm-diameter ROIs were placed at center of the field of view in Fiji to measure the mean fluorescence intensity. The 0 µM conditions were used for background subtraction. Mean intensities of the replicates were fit with a linear equation with the y-intercept constrained to zero. The resulting calibration curves-along with particle area and volume calculated from measured diameter- were used to calculate the concentration and number of loaded fluorescein or mCherry molecules.

## Acknowledgements

This work was supported by funding from the National Science Foundation to A.G.V. (Grant no. 1941966). We would like to thank L. Freddolino and M. E. DeSantis for their contributions to the discussion regarding cooperative cargo loading.

