## Supplemental Information for "Engineering modular cargo loading strategies for carboxysome-derived protein particles"

**Affiliations**

### Supplemental Figures

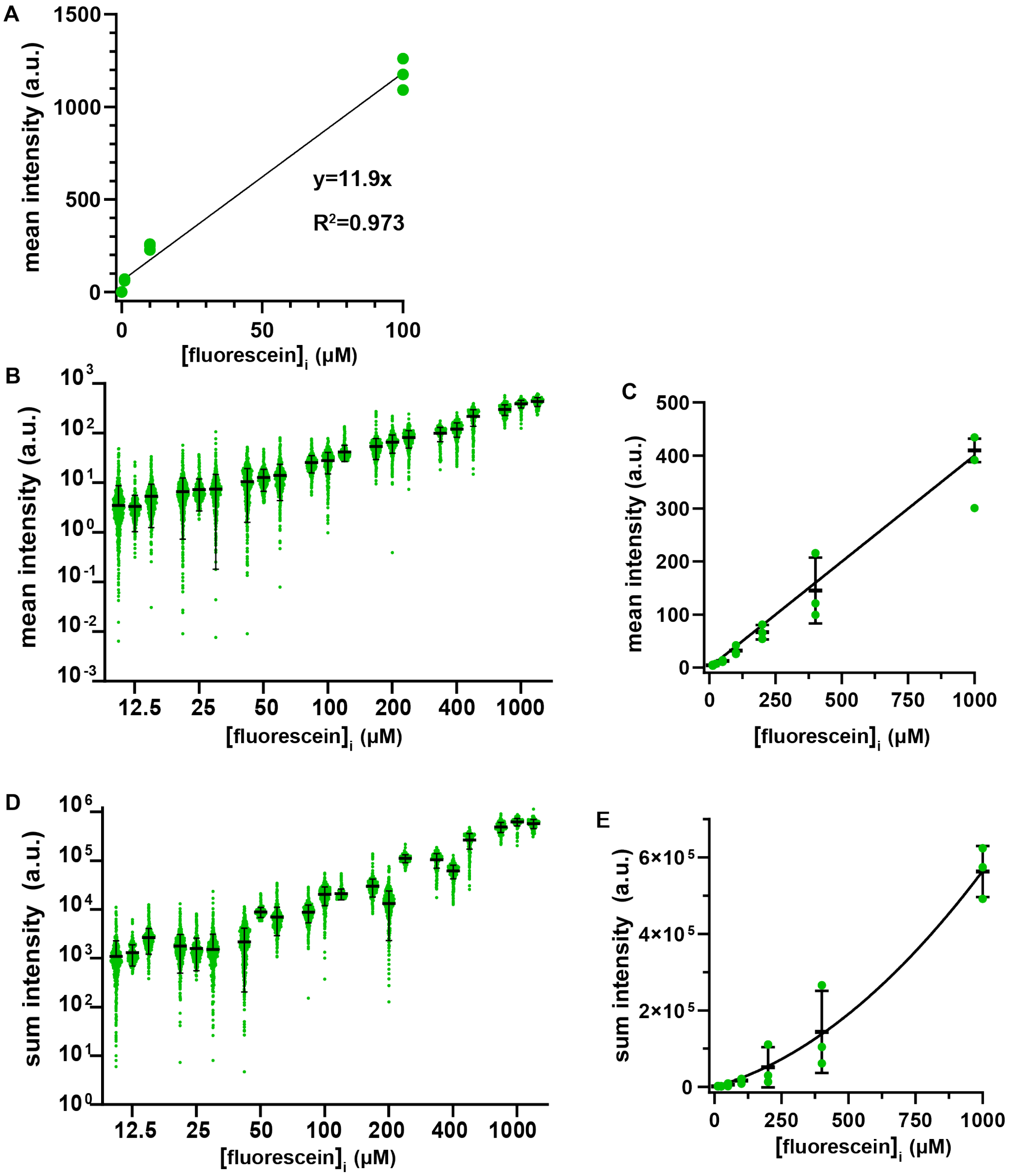


**Figure S1. Fluorescence intensity increases across fluorescein titrations. A.** Calibration curve of fluorescein. Green dots represent individual means across three replicates, and black bars indicate mean ± standard deviation (SD) for n = 3 technical replicates. Data were fit to a linear equation. **B-E.** Particle-associated fluorescence intensity of fluorescein. **(B)** Mean intensity at the indicated initial concentration of fluorescein. Green dots represent individual measurements across three replicates, and black bars indicate mean ± SD for n = 3 technical replicates. **(C)** Mean intensity as a function of initial fluorescein concentration. Green dots represent individual means across three replicates, and black bars indicate mean ± SD for n = 3 technical replicates. **(D)** Sum intensity of at the indicated initial concentration of fluorescein. Green dots represent individual measurements across three technical replicates. Black bars indicate mean ± SD for n = 3 replicates. **(E)** Sum intensity as a function of initial fluorescein concentration. Green dots represent means of replicates, and black bars represent overall mean ± SD for n = 3 technical replicates.

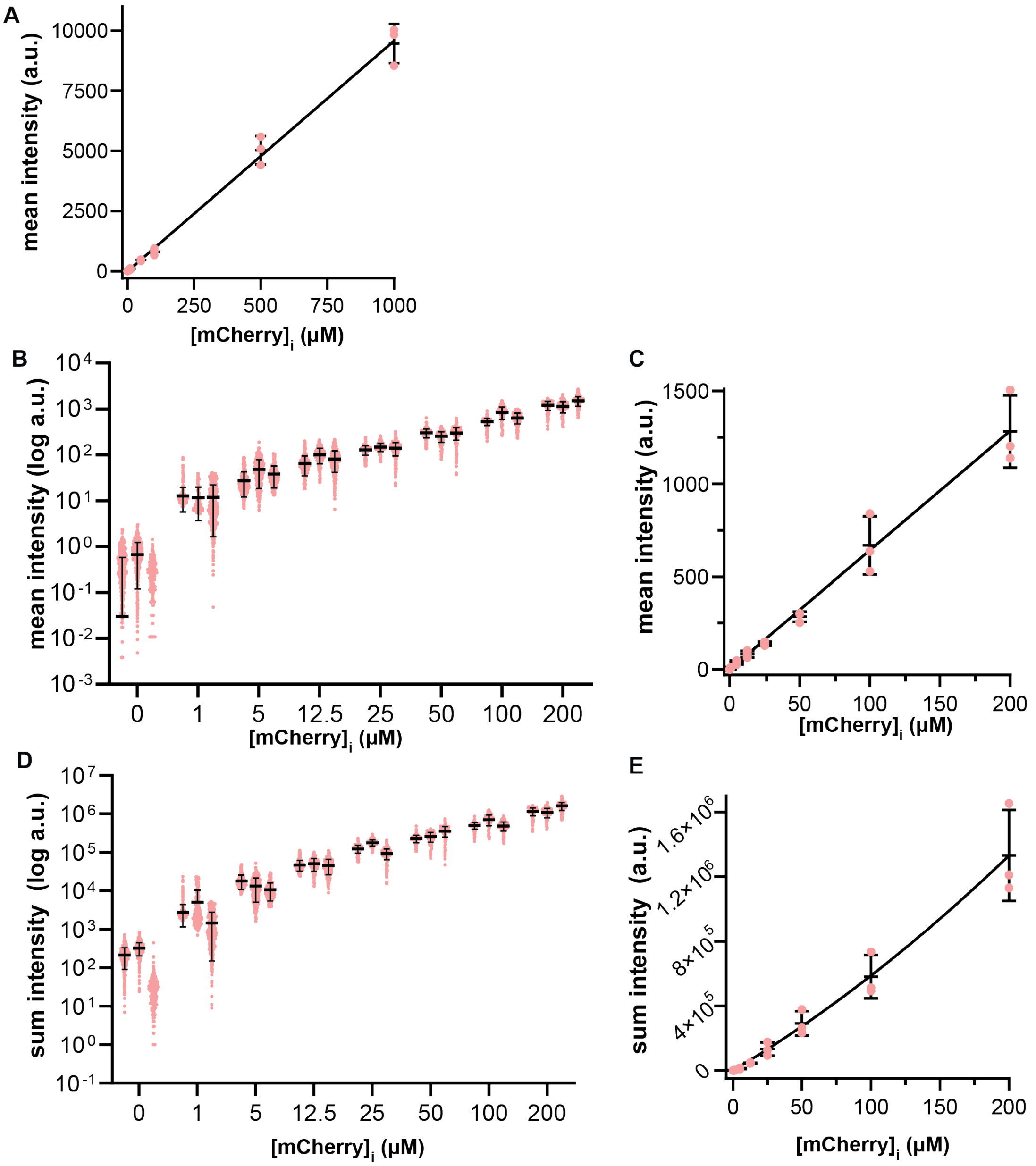


**Figure S2. Fluorescence intensity increases across mCherry titrations. A.** Calibration curve of mCherry. Light red dots represent individual means of technical replicates, and black bars indicate mean ± standard deviation (SD); n = 3. Data were fit to a linear equation. **B-E.** Particle-associated fluorescence intensity of mCherry. (B) Mean intensity at the indicated initial concentration of mCherry. Light red dots represent individual measurements for each technical replicate. Black bars indicate mean ± standard deviation (SD); n=3. (C) Mean intensity as a function of initial mCherry concentration. Light red dots represent means of technical replicates, and black bars represent overall mean ± SD; n = 3. (D) Sum intensity at the indicated initial concentration of mCherry. Light red dots represent individual measurements for each technical replicate. Black bars indicate mean ± SD; n=3. (E) Sum intensity of mCherry as a function of initial mCherry concentration. Light red dots represent means of technical replicates, and black bars represent overall mean ± SD; n=3. Data were fit to a quadratic equation.

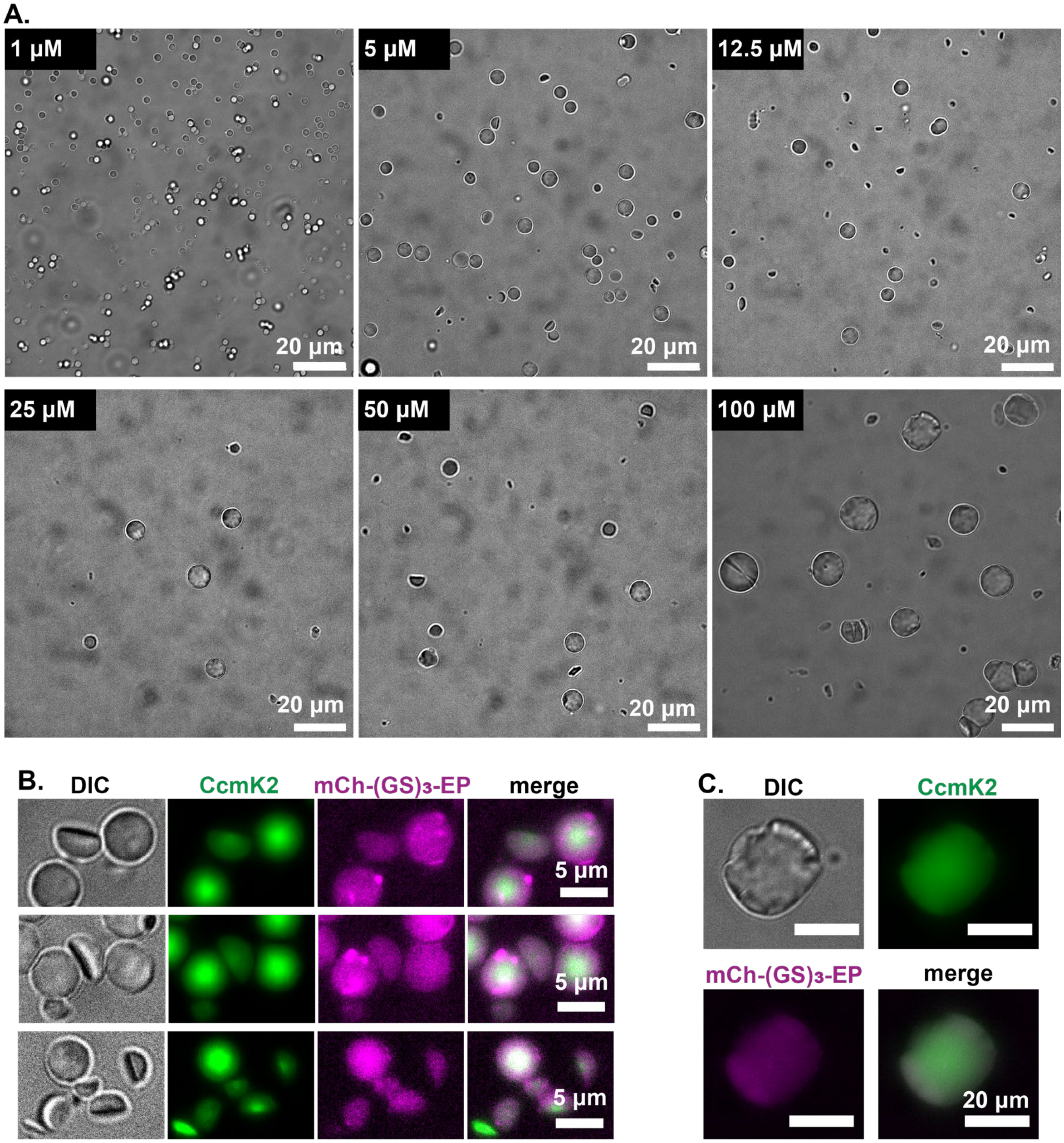


**Figure S3. mCh-(GS)$_3$-EP affects particle assembly. A.** Representative DIC fields of view of CcmK2 particles assembled at the indicated concentration of mCh-(GS)$_3$-EP. **B.** Representative DIC and fluorescence microscopy images of CcmK2 hemispheres. **C.** Representative DIC and fluorescence microscopy image of a CcmK2 particle with a wrinkly border.

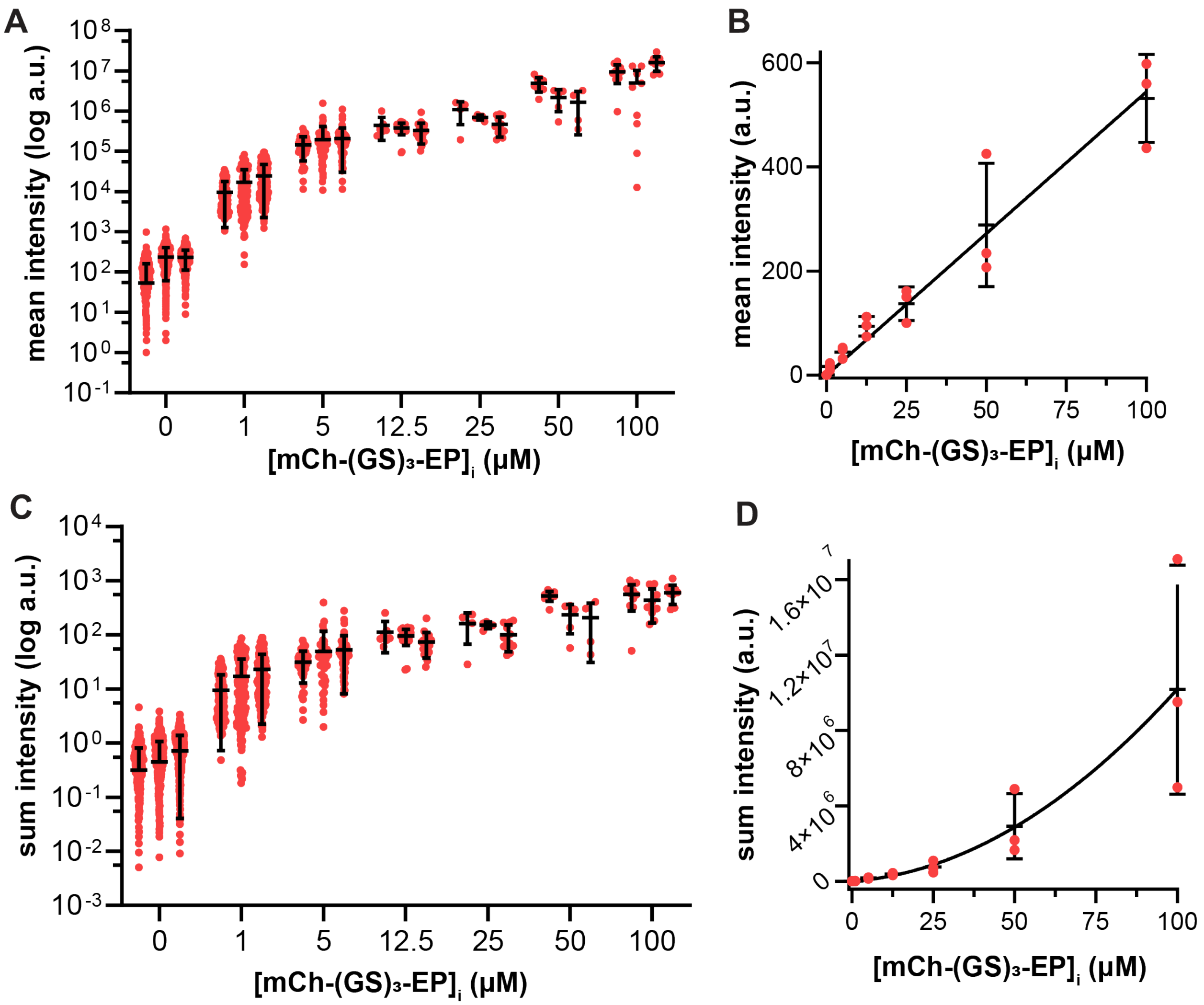


**Figure S4. Fluorescence intensity increases across mCh-(GS)3-EP titrations. A-D.** Particle-associated fluorescence intensity of mCh-(GS)3-EP. (A) Mean intensity at the indicated initial concentration of mCh-(GS)3-EP. Red dots represent individual measurements for each technical replicate. Black bars indicate mean ± standard deviation (SD); n = 3. (B) Mean intensity as a function of initial mCh-(GS)3-EP concentration. Red dots represent the means of technical replicates, and black bars represent overall mean ± SD; n = 3. Data were fit to a linear equation. (C) Sum intensity at the indicated initial concentration of mCh-(GS)3-EP. Red dots represent individual measurements for each technical replicate. Black bars indicate mean ± SD; n = 3. (D) Sum intensity as a function of initial mCh-(GS)3-EP concentration. Red dots represent means of technical replicates, and black bars indicate mean ± SD; n = 3. Data were fit to a quadratic equation.

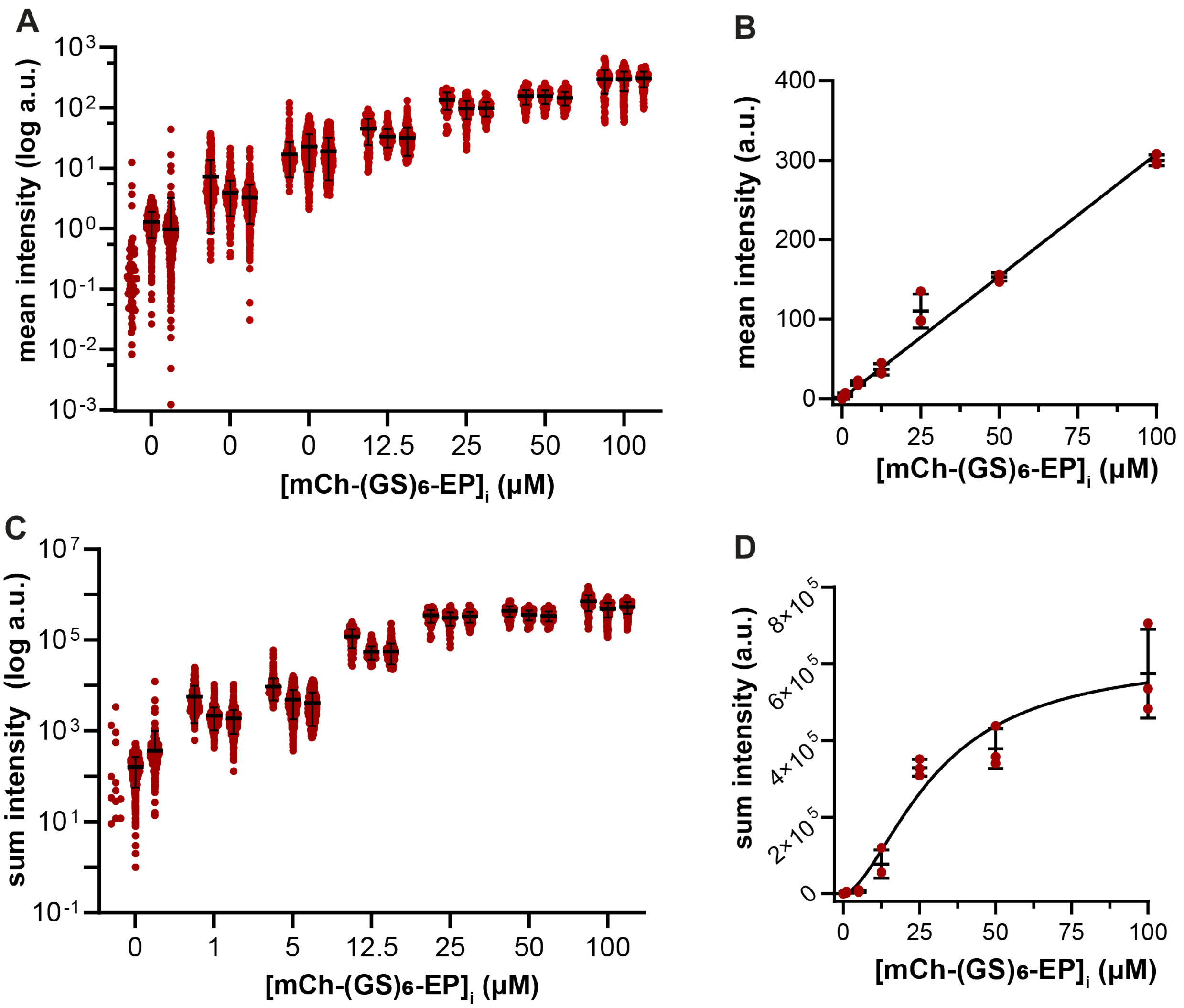


**Figure S5. Fluorescence intensity increases across mCh-(GS)$_6$-EP titrations. A-D.** Particle-associated fluorescence intensity of mCh-(GS)$_6$-EP. (A) Mean intensity at the indicated initial concentration of mCh-(GS)$_6$-EP. Dark red dots represent individual measurements for each technical replicate. Black bars indicate mean ± standard deviation (SD); n = 3. **B.** Mean intensity as a function of initial mCh-(GS)$_6$-EP concentration. Dark red dots represent the means of technical replicates, and black bars represent overall mean ± SD; n = 3. Data were fit to a linear equation. **C.** Sum intensity at the indicated initial concentration of mCh-(GS)$_6$-EP. Dark red dots represent individual measurements for each technical replicates. Black bars indicate mean ± SD; n = 3. **D.** Sum intensity as a function of initial mCh-(GS)$_6$-EP concentration. Dark red dots represent means of technical replicates, and black bars represent overall mean ± SD; n = 3. Data were fit to a quadratic equation.

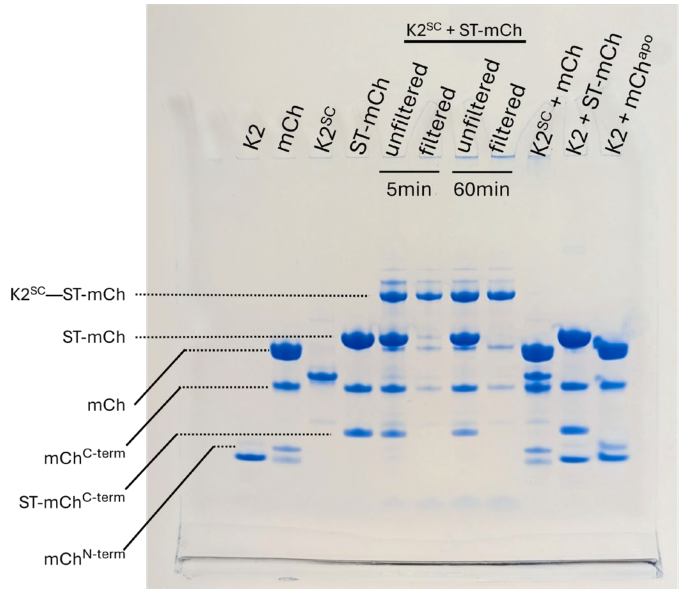


| construct | Abbreviated name | MW |
|---|---|---|
| CcmK2 | K2 | 11 kDa |
| His-mCherry | mCh | 28 kDa |
| His-mCh N-term trunc. | mCh$^{Nterm}$ | ~8 kDa |
| His-mCh C-term trunc. | mCh$^{Cterm}$ | ~20 kDa |
| His-SpyTag003-mCherry | ST-mCh | 30.9 kDa |
| His-ST-mCh N-term | ST-mCh$^{Nterm}$ | ~11 kDa |
| CcmK2[SpyCatcher003] | K2$^{SC}$ | 20.7 kDa |
| K2-SC—full length His-ST-mCh | K2$^{SC}$—ST-mCh | 51.6 kDa |
| K2-SC—His-ST-mCh N-term | K2$^{SC}$—ST-mCh$^{Nterm}$ | 31.2 kDa |

**Figure S6. CcmK2$^{SC}$ forms a complex with ST-mCherry.** *(left)* SDS-PAGE gel shows that CcmK2 with an internal SpyCatcher fusion (K2$^{SC}$) forms a complex with SpyTag fused to the N-terminus of mCherry (ST-mCh). *(right)* Table of all expected constructs and their molecular weights. The known molecular weights and typical distance traveled for CcmK2[17] (K2; Lane 1) and mCherry (mCh; Lane 2) were used to identify the remaining bands. mCh is known to form truncation products with an ~8 kDa N-terminal product and a ~20 kDa C-terminal product.[38] When ST-mCh is alone or in complex with K2$^{SC}$, the mCh-C-terminal band is the same as for mCh. Importantly, within 5 min, K2$^{SC}$ and ST-mCh complex formation is complete (unfiltered). Passing the reaction through a 100 kDa molecular-weight-cut-off spin filter removes unreacted ST-mCh and ST-mCh truncation products. The last three lanes show that complexes are not formed upon mixing K2$^{SC}$ and mCh, CcmK2 and ST-mCh, or and CcmK2 and mCherry.

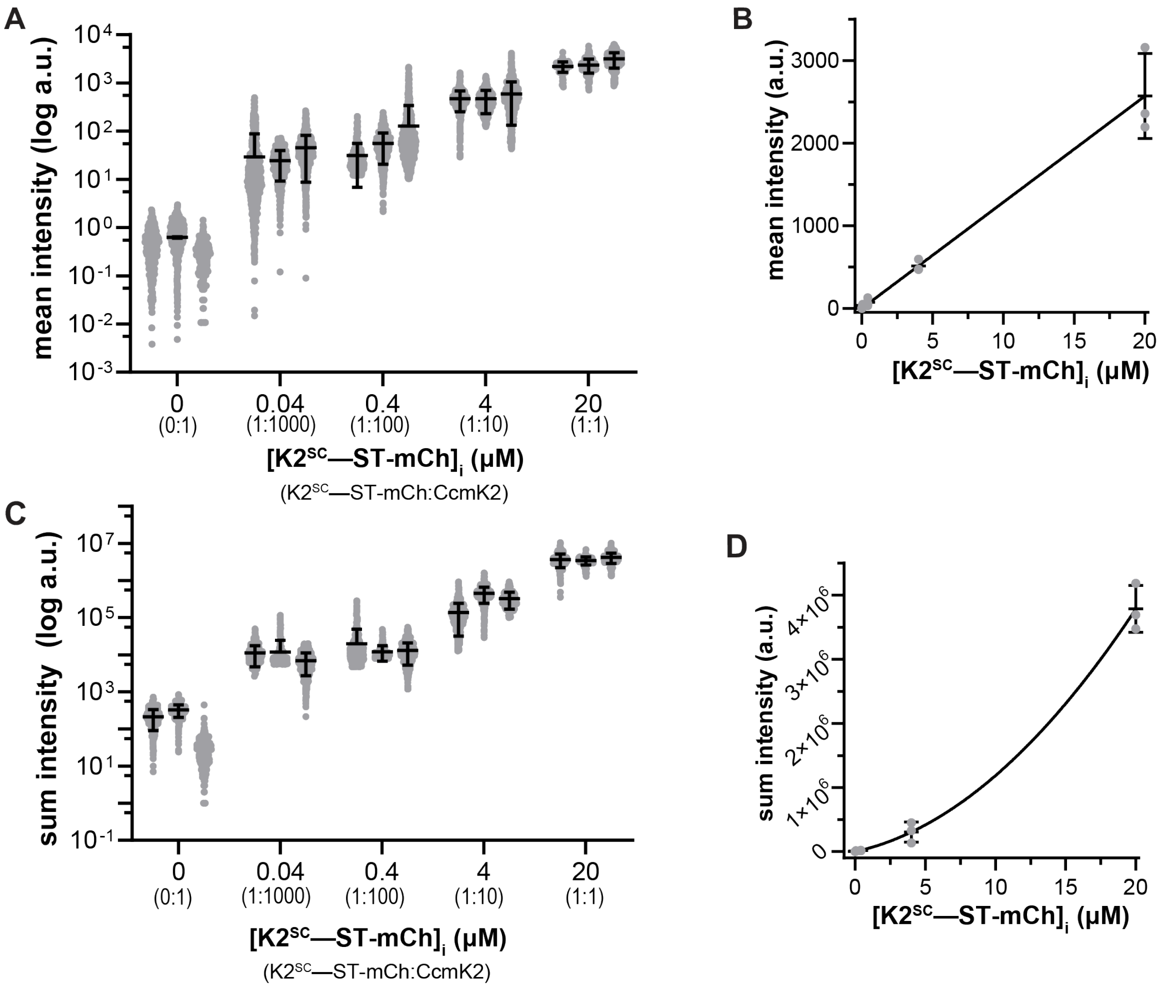


**Figure S7. Particle-associated fluorescence intensity increases across CcmK2$^{SC}$—ST-mCherry titrations. A-D.** Particle-associated fluorescence intensity of K2$^{SC}$—ST-mCh. (A) Mean intensity at the indicated initial concentration of K2$^{SC}$—ST-mCh. The molar ratio of K2$^{SC}$—ST-mCh:CcmK2 is also indicated. Gray dots represent individual measurements for each technical replicate. Black bars indicate mean ± standard deviation (SD); n = 3. **B.** Mean intensity as a function of initial CcmK2$^{SC}$—ST-mCh concentration. Gray dots represent the means of technical replicates, and black bars represent overall mean ± SD; n = 3. Data were fit to a linear equation. **C.** Sum intensity at the indicated initial concentration of K2$^{SC}$—ST-mCh. The molar ratio of K2$^{SC}$—ST-mCh:CcmK2 is also indicated. Gray dots represent individual measurements for each technical replicate. Black bars indicate mean ± SD; n = 3. **D.** Sum intensity as a function of initial CcmK2$^{SC}$—ST-mCh concentration. Gray dots represent means of technical replicates, and black bars represent overall mean ± SD; n = 3. Data were fit to a quadratic equation.
